# Targeting CBL ubiquitin ligase activation to downregulate tyrosine kinase signalling

**DOI:** 10.64898/2026.03.16.712190

**Authors:** Andrea J. Tench, Claire E. Martin, Craig D. Simpson, Leanne E. Wybenga-Groot, Diane Ly, Christopher Fladd, Melissa H. Elgie, Syed F. Ahmed, Roger Belizaire, Danny T. Huang, Anne-Claude Gingras, C. Jane McGlade

**Affiliations:** The Arthur and Sonia Labatt Brain Tumour Research Centre, 686 Bay St., Toronto, ON, M5G 0A42, Canada; Department of Medical Biophysics, University of Toronto, 101 College Street, Toronto, ON, M5G 1L7, Canada; Program in Cell and Systems Biology, 686 Bay St., Toronto, ON, M5G 0A42, Canada; Lunenfeld-Tanenbaum Research Institute, Mount Sinai Hospital, Sinai Health System, 600 University Ave, Toronto, ON, M5G 1X5, Canada; SPARC BioCentre, The Hospital for Sick Children, 686 Bay St., Toronto, ON, M5G 0A42, Canada; Dana-Farber Cancer Institute, 75 Francis St, Boston, MA, 02115, USA; Cancer Research UK Scotland Institute, Garscube Estate, Switchback Road, Glasgow G61 1BD, UK; School of Cancer Sciences, University of Glasgow, Glasgow G61 1QH, UK; Department of Molecular Genetics, University of Toronto, 1 King’s College Circle, Toronto, Ontario, M5S 3K3, Canada

## Abstract

The CBL E3 ubiquitin ligase is a critical regulator of tyrosine kinase (TK) signalling. CBL activity is regulated by a feedback loop in which an active TK phosphorylates CBL, relieving its autoinhibited conformation, and allowing ubiquitination of substrates, including TKs, leading to their degradation. Binding of the Src Like Adapter Protein 2 (SLAP2) to CBL can also activate autoinhibited CBL and promote substrate ubiquitination. Using an engineered CBL mutant in the SLAP2 binding interface, termed RE CBL, that mimics activation by SLAP2 binding, we characterized the cellular functions of this CBL activation mechanism. Comparison of wildtype and RE CBL interactomes using MiniTurboID showed extensive and overlapping interaction networks, with a discrete subset of proteins, including the known substrate, epidermal growth factor receptor (EGFR), as well as endocytic factors such as EPS15, in higher abundance with RE CBL compared to wildtype. Consistent with these observations, RE CBL interacted more readily with EGFR, enhanced EGFR internalization, and attenuated downstream signalling compared to WT. In *Cbl* null hematopoietic cells, RE CBL expression reduced sensitivity to cytokines IL-3 and GM-CSF, and decreased activation of the Src-family kinase Lyn. Furthermore, we optimized and conducted a small molecule screen to identify a group of structurally related compounds that, like SLAP2 binding, promoted CBL activation *in vitro*. Together these findings provide proof of concept for targeting CBL activity to downregulate TK signalling.

## Introduction

The post-translational attachment of the small protein ubiquitin to protein substrates can direct multiple outcomes, including protein degradation via the proteasomal and lysosomal pathways.^1,2^ The ubiquitin cascade in the human proteome includes two known ubiquitin E1 activating enzymes, approximately 40 E2 conjugating enzymes, and over 600 different E3 ubiquitin ligases that determine substrate specificity for ubiquitination.^3^ CBL is an E3 ubiquitin ligase that negatively regulates receptor and non-receptor tyrosine kinases (TK) that mediate signalling pathways controlling cellular growth, proliferation, migration, amongst other processes.^4,5^ Overactivation of TK pathways is a common oncogenic driver, such that CBL typically acts in a tumour suppressive function to promote TK downregulation.^6,7^

CBL is composed of a tyrosine kinase binding domain (TKBD), linker-helix region (LHR) and E2 binding RING domain, followed by protein-protein interaction motifs in the carboxy terminus, including a proline rich region and tyrosine residues that can be phosphorylated and serve as docking sites for additional signalling proteins.^8–10^ Structural studies have shown that inactive CBL exists in an autoinhibited conformation with the TKBD and LHR region in close association, and the RING domain oriented away from the substrate binding region of the TKBD.^11^ Phosphorylation within the LHR at Tyr371, disrupts the TKBD-LHR interface resulting in a conformational change that reorients the RING domain, placing the E2 active site in close proximity to TKBD bound substrate to facilitate ubiquitin transfer.^11^ CBL-mediated ubiquitination promotes receptor internalization and trafficking to lysosomes for degradation, as well as targeting cytoplasmic signalling proteins for degradation through direction to the proteasome.^4^ In addition to its role as an E3 ubiquitin ligase, the carboxy terminal SH2 and SH3 domain interaction motifs facilitate CBL function as an adaptor protein, helping to nucleate proteins at receptor complexes involved in signal transduction and endocytic events.^8^

Mutations in *CBL* that abrogate its ubiquitin ligase activity have been identified in subsets of myeloid leukemias. *CBL* mutations occur in approximately 5% of myeloid leukemias, including 15-20% of Juvenile Myelomonocytic Leukemia (JMML) and 15% of Chronic Myelomonocytic Leukemia (CMML) cases.^12–16^ The most frequent *CBL* mutations include missense mutations at Tyr371 within the LHR and the zinc-coordinating residues of the RING domain, both of which disrupt CBL E3 activity.^17,18^ While the tumour suppressive E3 ligase function is lost, CBL mutant proteins can also promote oncogenic signalling through retained adaptor function, in which the intact TKBD allows sustained interactions with receptor complexes.^19–23^ For example, mutant CBL has been shown to have increased associations with the granulocyte-macrophage colony-stimulating factor (GM-CSF) receptor β common (βc) and GM-CSF specific α (GMRα) chain, as well as downstream signalling proteins including the Src family kinase Lyn and the p85 subunit of PI3K.^19–21^ Indeed, a characteristic feature of JMML and CMML is hypersensitivity to GM-CSF, and this has been shown experimentally to occur in part due to increased PI3K signalling mediated by Lyn.^17,20,21,24^

Dysregulation of CBL activity has also been observed in several solid tumours through changes in its phosphorylation status and decoupling from substrates through substrate alterations or protein-protein interactions that sequester CBL from its target.^25–31^ For instance, oncogenic mutations in the epidermal growth factor receptor (EGFR) that disrupt CBL binding are common in tumours such as non-small cell lung cancer (NSCLC), glioblastoma, colorectal, breast, and ovarian tumours.^32–34^ Additionally, multiple protein-protein interactions that prevent CBL from binding and ubiquitinating EGFR and other TKs are altered in tumours.^27–31,35–38^

CBL is recruited to active receptor complexes directly via the TKBD or indirectly by other adaptor proteins such as GRB2 and SLAP2. For example, the amino-terminal SH3 domain of GRB2 interacts with the proline rich region of CBL while the GRB2 SH2 domain binds active receptor tyrosine kinases like the EGFR.^39,40^ In hematopoietic cells, the Src-like adaptor proteins SLAP and SLAP2 are required for CBL-mediated regulation of antigen receptors including B and T cell receptor components, as well as downregulation of growth factor and cytokine receptors such as GM-CSFR.^41–47^ In addition to recruiting CBL to activated TKs, SLAP2 binding also promotes CBL ubiquitination activity, independent of tyrosine phosphorylation.^48^ Structural characterization revealed that SLAP2 binds a unique pocket of the CBL TKBD, overlapping the region where the linker helix region resides in the closed, autoinhibited conformation.^48,49^ Here we characterize a mutant form of CBL (RE CBL) as a model for SLAP2-mediated CBL activation and potential mechanism to be targeted by small molecule modulation of CBL. We determine the RE CBL mutant interactome and its effects on signalling downstream of EGFR and GM-CSFR. We further optimized and conducted a small molecule screen to identify compounds that promote CBL activity *in vitro*, establishing tool compounds for future phenotypic exploration of promoting CBL activation.

## Results

### SLAP2 binding site mutations activate CBL E3 ligase activity

The interaction between SLAP2 and CBL involves an α-helix in the carboxy-terminus of SLAP2 (residues 237-255) that binds in a cleft of the CBL TKBD opposite to the pTyr-peptide binding region (**Figure 1a**).^48^ In the autoinhibited form of CBL, this region is typically occupied by the LHR, suggesting that SLAP2 binding could disrupt the autoinhibited CBL conformation (**Figure 1b**). In agreement, the RE CBL mutant that contains amino acid substitutions at Ala223 to Arg (A223R) and Ser226 to Glu (S226E) in this region of the TKBD (**Figure 1b**), disrupted the interaction with SLAP2, and also promoted CBL *in vitro* ubiquitin ligase activity (**Figure 1c**).^48^ Since oncogenic *CBL* mutations in the LHR, such as those at Tyr371, also disrupt the LHR-TKBD interface, we tested the potential transforming activity of RE CBL in a focus forming assay. Similar to wildtype (WT) CBL, expression of RE CBL did not cause foci formation in NIH3T3 cells, in contrast to oncogenic CBL variants such as Y371H, Y371C, Y371S and 70Z CBL (**Figure 1d-f**). These findings suggest that disruption of the TKBD-SLAP2 binding interface can activate CBL E3 ligase function *in vitro*, without causing transforming activity in cells.

**Figure 1:**
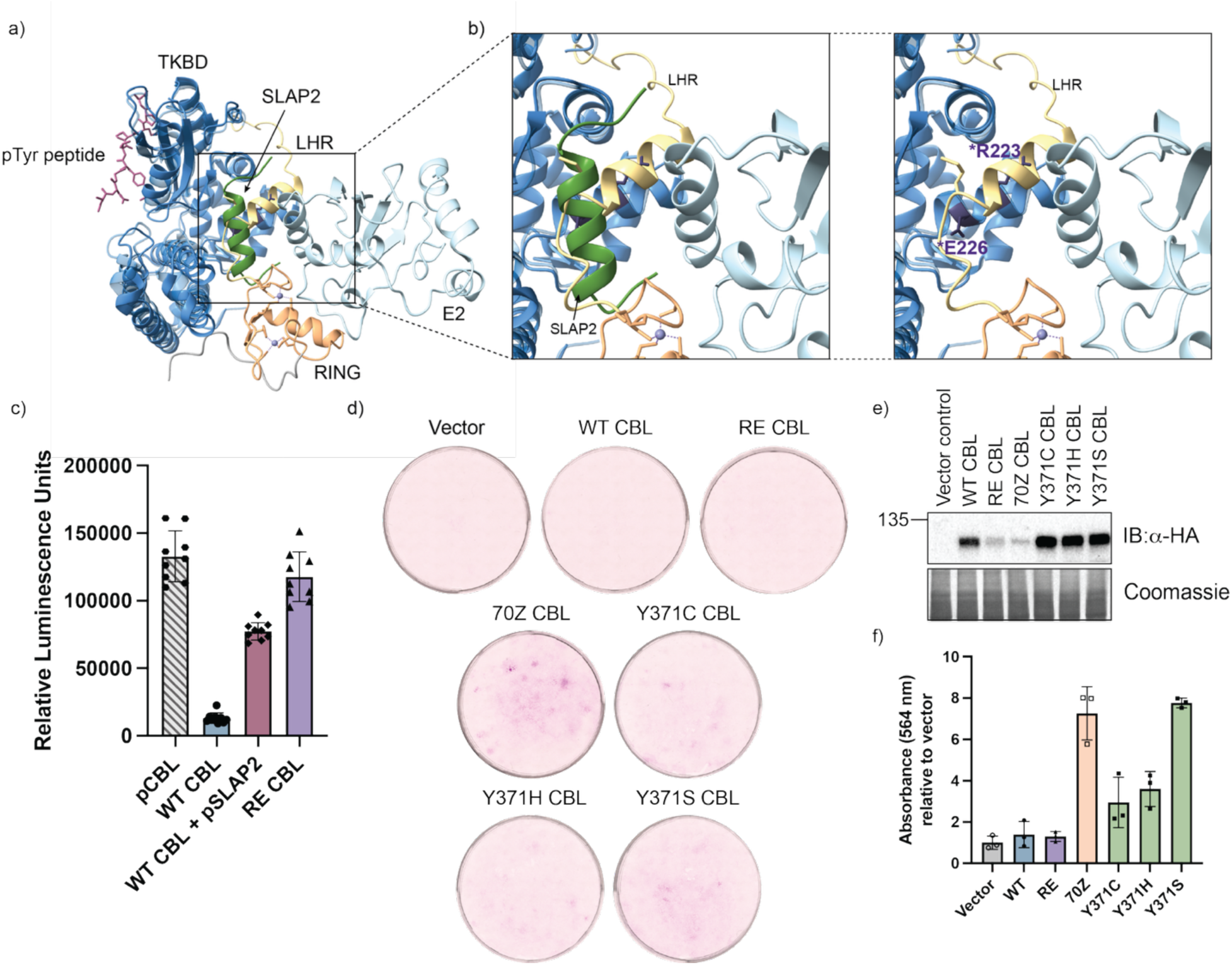
RE CBL mutation mimics SLAP2-mediated activation of autoinhibited CBL. **a)** CBL tyrosine kinase binding domain (TKBD) in dark blue associated with SLAP2 tail helix in green (PDB: 6XAR) overlayed with TKBD-linker helix region (LHR)-RING (light blue, yellow and orange respectively), associated with a pTyr peptide (pink) modeling substrate binding and a bound E2 (PDB: 1FBV). **b)** Left inset shows the overlay of the LHR and SLAP2 in the autoinhibited conformation. Right inset shows the A223R and S226E mutations modeled in purple. **c)** *In vitro* activity of phosphorylated CBL (pCBL - diluted), wildtype (WT) CBL, WT CBL with pSLAP2 in the reaction, and RE CBL as assessed by E3Lite assay. **d)** NIH3T3 focus forming assay of cells transduced with vector control or indicated CBL form expression vectors. **e)** Blot shows expression of the HA-tagged CBL proteins. **f)** Bar plot shows the dye absorbance read at 564 nm.

### Proximity interactomes of CBL and mutant RE CBL

Proximity dependent biotinylation coupled with mass spectrometry (PDB-MS) with MiniTurboID was conducted to compare the interactomes of WT and mutant RE CBL.^50^ HeLa Flp-IN T-REx cells were generated with inducible expression of WT or RE CBL with carboxy-terminal tags with MiniTurbo-FLAG, as well as a control cell lines expressing the FLAG-tagged MiniTurbo protein (**Supplementary Figure 1a**). Immunofluorescence confirmed that biotinylated proteins localized with CBL-MiniTurbo-fusion proteins (**Supplementary Figure 1b**) and that CBL-MiniTurbo fusions co-localized with EGFR in response to EGF treatment (**Supplementary Figure 1c**). MiniTurbo WT and RE CBL expressing HeLa cells were labeled with biotin over a two hour period with or without EGF treatment, lysed, and biotinylated proteins were isolated and identified by mass spectrometry. A total of 237 unique CBL proximity interactors with a BFDR of 0.01 or below were identified across all conditions. Gene ontology analysis and literature search of the interactors revealed involvement in signal transduction, endocytosis, trafficking and intracellular transport, cytoskeletal, adhesion and migration, cell cycle regulation, and DNA and RNA processing (**Figure 2a**). Most proteins were identified in all conditions albeit with varying abundance and significance scores as described below (**Supplementary Figure 2**).

**Figure 2:**
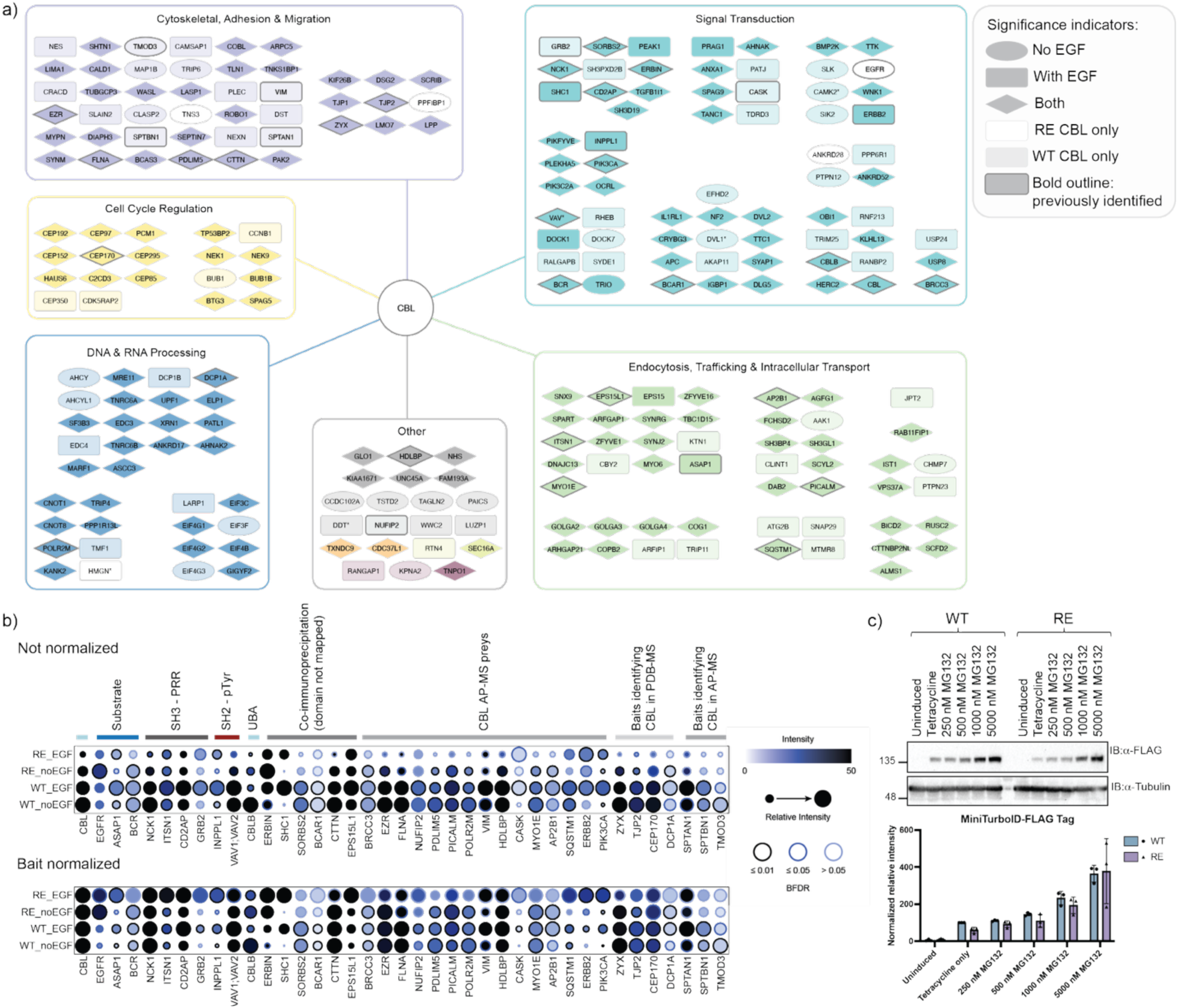
MiniTurboID interactome of wildtype and RE CBL with and without EGF stimulation. **a)** Summary figure of significant interactors from the proximity dependent biotinylation in HeLa Flp-In TREx cells. Data was acquired and processed from 3 independent biological replicate experiments. Interactors were scored as significant based on a BFDR of 0.01 or less and g:Profiler was used to make categorizations and confirmed through searching Uniprot and Gene Cards annotations. Interactors significant only with no EGF treatment are shown as ovals, only with EGF are shown as rounded squares, or in both conditions are shown as diamonds. Interactors significant with wildtype CBL only have lighter shading. Interactors significant with RE CBL only have white nodes. Known CBL interactors from BioGrid and literature search are indicated with a bold outline (see legend on right). **b)** Dot plot to compare known CBL interactors across conditions. Significant interactors (BFDR σ; 0.01 in any condition) were searched in BioGrid and in the literature for known association with CBL. The upper dot plot is representative of unnormalized data, while the lower dot plot shows the data normalized to the bait count to account for the lower CBL levels with RE CBL. Dot size is indicative of relative abundance, and the outline indicates the BFDR score per the legend on the right. **c)** Protein lysates from HeLa Flp-In T-REx cells expressing C terminally tagged FLAG-MiniTurboID WT or RE CBL treated with the indicated concentrations of MG132 for 16 hours were resolved by SDS-PAGE and immunoblotted with anti-FLAG (CBL) and anti-tubulin as a loading control. Bands were quantified and FLAG signals were normalized to tubulin and expressed as a percentage of the WT CBL induced (tetracycline only) signal. Plot shows the mean and individual normalized values from three independent biological replicates with error bars indicating the standard deviation.

In the identified proximity interactome, 39 proteins have been previously characterized as CBL interactors (**Figure 2b**). These proteins included CBL substrates, EGFR, ASAP1 and BCR.^51–55^ In addition, NCK1, ITSN1, CD2AP and GRB2 were previously shown to bind the proline rich region (PRR) of CBL through SH3 domain mediated interactions.^56–59^ Additionally, INPPL1 (SHIP2) and VAV1 are known to interact with carboxy-terminal phosphorylated tyrosines of CBL through SH2 domain mediated interactions.^60,61^ CBL and CBL-b are also known to homo- and hetero-dimerize through the ubiquitin association (UBA) domain at the carboxy terminus.^62,63^ Additionally, ERBIN, SHC1, SORBS2, BCAR1, CTTN and EPS15L1 have been experimentally shown to interact with CBL, though their interactions have not been mapped to particular domains.^30,52,64–67^ Additional preys were also identified in published CBL affinity purification mass spectrometry (AP-MS) experiments.^52,68,69^ As well, baits that identified CBL as a prey in PDB-MS and AP-MS experiments were also captured.^70–72^ Notably, normalizing the intensity values to the CBL levels to account for bait differences suggests similar abundance of many preys identified by WT and RE CBL (**Figure 2b**). The high number of shared interactors between WT and RE CBL, including known CBL binding partners and proteins involved in expected processes, demonstrates that the RE mutation in the TKBD does not disrupt CBL protein interactions or localization.

A trend toward lower intensity values for RE CBL proximity interactors compared to WT CBL, especially when cells were treated with EGF, was observed (**Supplementary Figure 3a**). The reduced intensity values for the RE CBL bait likely contributed to decreased prey intensity overall. We observed equalized levels of RE and WT CBL following treatment with increasing concentrations of a proteasomal inhibitor, MG132, consistent with reduced RE CBL expression due to enhanced activation leading to autoubiquitination and degradation (**Figure 2c; Supplementary Figure 3b**).

### Changes in the CBL interactome in response to EGF stimulation suggest altered EGFR signalling and endocytic dynamics in the presence of RE CBL

To delineate the effect of EGF treatment on CBL interacting partners and whether there were unique responses between WT and RE CBL, we compared the set of interactors that scored as significant for both proteins. A core group of interactors that increased in abundance in response to EGF treatment with either WT (**Figure 3a**) or RE CBL (**Figure 3b**) included PRAG1, PEAK1, SHC1, INPPL1, ASAP1, MYO6, ZFYVE16, PIK3CA, and DOCK1. In WT CBL expressing cells, CASK and ERBB2 also significantly increased, while GRB2 and USP8 also trend toward increasing with EGF treatment (**Figure 3a**). In RE CBL expressing cells, EPS15 was notably increased in abundance following EGF stimulation compared to WT CBL (**Figure 3b**). Many of these proteins have known interactions and involvement in EGFR signalling. In particular, most proteins that increased in abundance following EGF treatment were adaptor and scaffold proteins, in addition to enzymes involved in signal transduction (**Figure 3c**).

**Figure 3:**
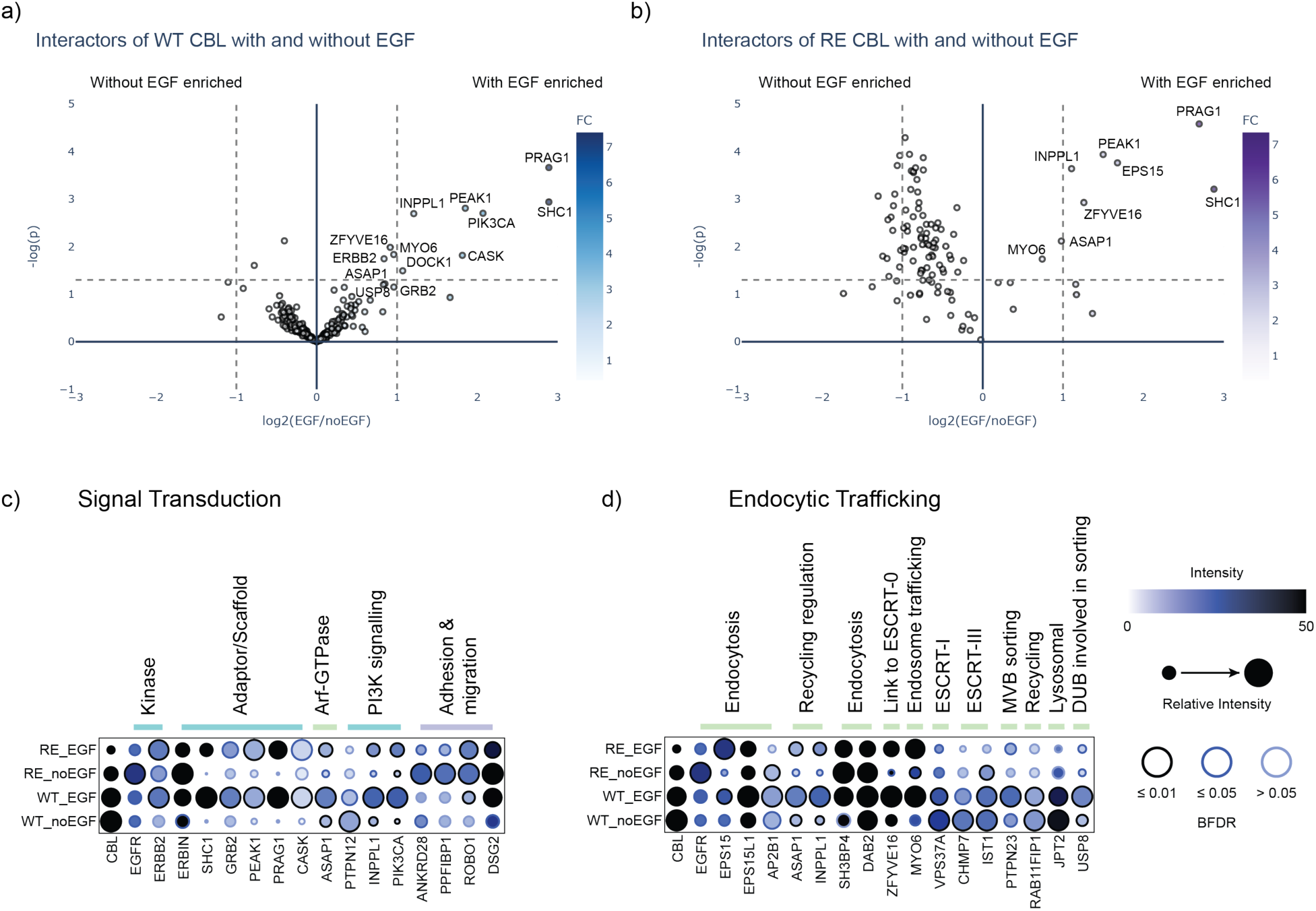
Comparison of WT and RE CBL proximity interactors in response to EGF treatment. **a)** Interactors with a BFDR of 0.01 and below for WT CBL without or with EGF treatment are plotted. Triplicate spectral counts in no EGF compared to EGF treated conditions were analyzed by T test with the -log of the p value plotted on the y-axis. The log2 of the ratio of EGF to no EGF treatment is plotted on the x-axis. Vertical lines indicate a fold change of 2 and over in either direction. The horizontal dotted line indicates a p value of 0.05, points above this line are interactors with a p value scoring less than 0.05. The colouring of the dots is relative to the ratio of EGF treated to no EGF per the scale on the right. **b)** Shows the same analysis as in **a)** for RE CBL interactome data without or with EGF treatment. Dotplots of interactors of interest involved in **c)** signal transduction and **d)** endocytic trafficking are shown comparing relative abundance of the interactors across the four conditions indicated with the abundance and BFDR score outlined in the legend to the right.

Despite the overall lower abundance of proteins identified with RE CBL, several proteins including EGFR, SH3BP4 and EPS15 were enriched compared to WT CBL in either condition (**Figure 3d)**. SH3BP4 is involved in endocytic processes through promoting incorporation of target proteins, such as α5-integrin, in clathrin coated pits and is recruited at least in part by EPS15.^73,74^ EPS15 is recruited to active EGFR through its ubiquitin interaction motif, and coordinates interactions with endocytic adaptor proteins in a ubiquitin dependent manner, facilitating internalization and trafficking of the receptor.^75–79^ Other endocytic proteins were similarly identified between WT and RE CBL and proteins involved in later endosomal trafficking were lower in abundance in the RE CBL interactome (**Figure 3d**). Together, the subset of uniquely elevated interactors with RE CBL point to a potential change in EGFR engagement and early endocytic dynamics in the presence of RE CBL.

### Enhanced EGFR binding and phosphorylation of RE CBL

A key finding in the MiniTurboID dataset was that EGFR was more abundant in the RE CBL condition compared to WT CBL (**Figure 3d**). To determine how the RE mutation impacted the association between EGFR and CBL, HeLa cells with inducible expression of carboxy terminal fusions of a 3XFLAG tag to WT or RE CBL cells were treated with increasing concentrations of EGF for two minute stimulations, to capture the initial recruitment of CBL to EGFR. Although RE CBL expression is lower than WT due to autoubiquitination (**Figure 4d; Supplementary Figure 3b**), FLAG-RE CBL immunoprecipitates captured significantly greater levels of EGFR, relative to the CBL levels, from cells treated with 50 and 100 ng/mL EGF (**Figure 4a**). Similarly, greater relative FLAG-RE CBL was co-immunoprecipitated with EGFR, most notably at 5 ng/mL EGF (**Figure 4b)**. The observed increase in relative RE CBL and EGFR association by co-immunoprecipitation is consistent with the trend observed in MiniTurboID experiment and indicates that RE CBL is more efficiently recruited to activated EGFR.

**Figure 4:**
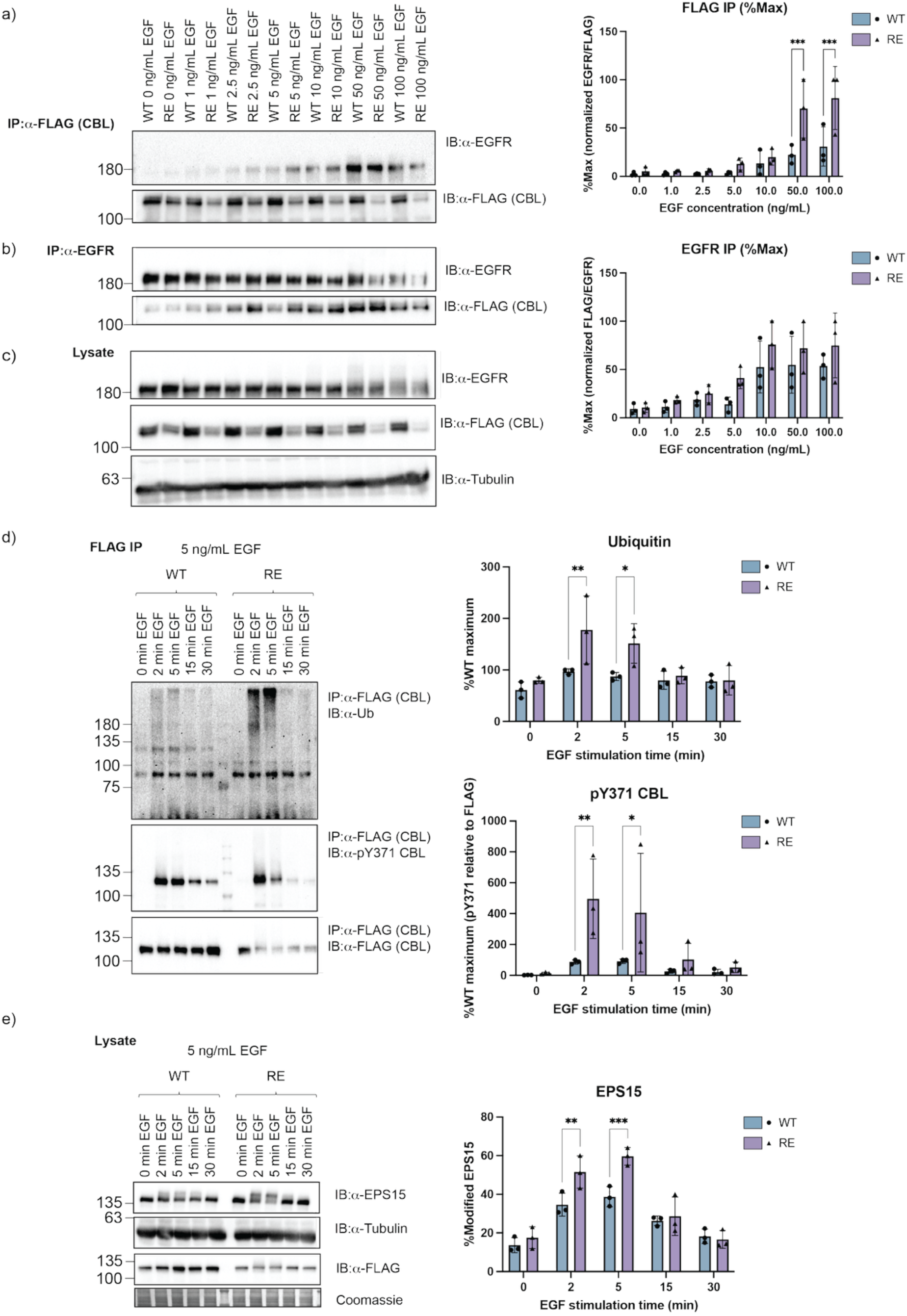
RE CBL promotes EGFR engagement, CBL activation and EPS15 modification. HeLa Flp-In T-REx cells induced to express carboxy terminally 3XFLAG-tagged WT or RE CBL and treated with increasing concentrations of epidermal growth factor (EGF) for 2 minutes. Lysates immunoprecipitated (IP) with **a)** anti- FLAG (top) and **b)** EGFR (middle) and immunoblotted for EGFR and FLAG (CBL) from the same membrane. For **a)** and **b)** quantification of immunoblots are shown to the right. EGFR signals were normalized to FLAG for the FLAG IP (upper) and FLAG signals normalized to EGFR for the EGFR IP (lower). Results normalized as the %Maximum signal from three independent biological replicates are plotted. **c)** Lysates were also immunoblotted for total EGFR, total FLAG (CBL) and tubulin levels (bottom). **d)** HeLa Flp-In T-REx cells expressing carboxy terminally 3XFLAG-tagged WT or RE CBL were induced, serum starved then treated with 5 ng/mL EGF for the indicated times. Lysates were immunoprecipitated with anti-FLAG (CBL), and samples were divided into three immunoblots using anti-ubiquitin, anti-pY371 CBL and anti-FLAG (CBL). Ubiquitin signal (upper graph) and pY371 relative to FLAG (CBL) levels (lower graph) were quantified from three independent experiments and expressed as a percentage of the maximum signal with WT CBL. **e)** Corresponding lysates from the immunoprecipitations in **d)** were immunoblotted with anti-FLAG (CBL) and membranes were stained with Coomassie as loading control, or anti-EPS15 with a corresponding tubulin blot. The upper EPS15 band (modified) was quantified relative to total EPS15 levels from three independent experiments and expressed as the percentage of total EPS15. For all plots, the bars represent the mean while the individual points from each experiment are also shown. Error bars indicate the standard deviation. Data were analyzed by two-way ANOVA with Tukey’s correction (* indicates p < 0.05, ** p < 0.002 and *** p < 0.0002).

To further characterize the impact of the more efficient association of EGFR and RE CBL, we investigated the status of CBL phosphorylation and ubiquitination through immunoprecipitation. At lower EGF concentrations (5 ng/mL), we observed that RE CBL is more readily phosphorylated at Tyr371 and associated with increased ubiquitination activity in CBL (FLAG) immunoprecipitations, particularly at early time points of stimulation (**Figure 4d**). Further, in agreement with the increased EPS15 abundance observed in the RE interactome, we also observed more rapid modification of EPS15 following EGF stimulation in the presence of RE CBL, indicated by the higher molecular weight band observed, which likely corresponds to ubiquitinated EPS15 (**Figure 4e**).^80^ This coincides with more rapid recruitment of EPS15 to EGFR, which has been shown to be mediated by CBL activity.^76^ Together these results support findings in the MiniTurboID dataset and indicate that RE CBL more readily engages substrate (EGFR), leading to its more efficient Tyr371 phosphorylation and increased ubiquitination activity.

### RE CBL enhances EGFR internalization and attenuation of downstream signalling

To investigate the functional consequences of RE CBL activity on EGFR downregulation, we determined the effects of RE CBL expression on EGFR internalization by surface biotinylation, and isolation of surface and total EGFR over time course experiments. EGFR surface levels significantly decreased in RE CBL expressing cells compared to WT CBL expressing cells at low (5 ng/mL) EGF concentrations (**Figure 5a,b**) but not at a higher (50 ng/mL) EGF concentration (**Supplementary Figure 4**). Overexpression of both WT or RE CBL promoted more rapid loss of cell surface EGFR compared to FLAG control expressing cells, demonstrating that the observed effect was dependent on the overexpressed CBL. While total EGFR levels also decreased following stimulation, there was no significant difference in the rate of total EGFR loss between WT and RE CBL expressing cells (**Figure 5b**, **Supplementary Figure 4**).

**Figure 5:**
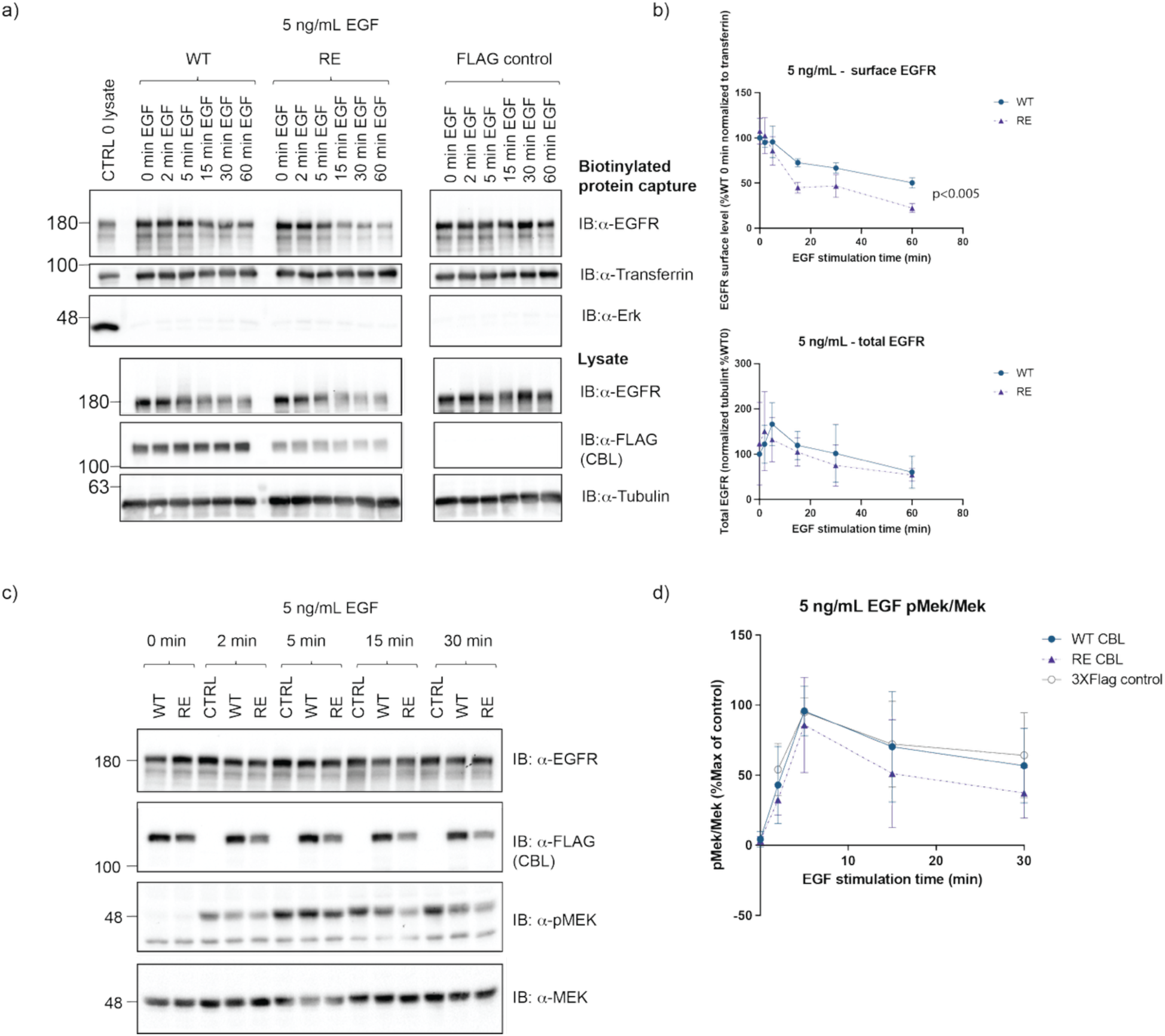
RE CBL promotes EGFR internalization and signal attenuation. **a)** HeLa cells induced to express WT or RE CBL, or FLAG control, were treated with 5 ng/mL EGF for the indicated times. Cell surface proteins were biotinylated and captured from cell lysate on streptavidin agarose and analyzed by immunoblotting with anti-EGFR, anti-transferrin as a positive control for surface proteins, and negative control anti-Erk. Figures show representative blots from independent triplicate experiments. **b)** Quantification of EGFR and transferrin immunoblots. Surface biotinylated EGFR levels (upper graph) were normalized to the transferrin level and then to the signal for WT CBL at 0 ng/mL EGF (for both WT and RE). Total EGFR levels (lower graph) were normalized to tubulin level and then to the signal for WT CBL at 0 ng/mL EGF (for both WT and RE). The mean and standard deviation from three independent experiments are shown. **c)** Protein lysates from cells treated with 5 ng/mL EGF for the indicated times were immunoblotted with anti-EGFR, anti-phosphoMEK(pSer217/221) or anti-MEK1/2. **d)** Quantification of phospho-MEK levels normalized to total MEK levels from three biological replicates is shown as a percentage of maximum signal in the control cell line. Data were analyzed by two-way ANOVA with Tukey’s correction (significance indicated on plot).

In agreement with enhanced downregulation of surface EGFR, in the presence of RE CBL we also observed a trend in more efficient attenuation of activated MEK at 5 ng/mL EGF (**Figure 5c**) but not 50 ng/mL EGF treatment (**Supplementary Figure 5b,d**). Differences in downregulation of activated Erk or Akt were not observed in these experiments (**Supplementary Figure 5**). Together these results suggest that RE CBL promotes recruitment of endocytic adaptors such as EPS15, more efficient EGFR internalization, and attenuates downstream signalling.

### RE CBL attenuates cytokine-dependent proliferation and Lyn activation

To compare the effects of RE CBL expression to WT CBL and the oncogenic Y371H (YH) CBL mutant on cytokine-dependent proliferation and signalling, we generated doxycycline inducible cell lines using the cytokine-dependent murine myeloblast 32D cell line with endogenous *Cbl* knocked out.^21^ Expression of Y371H CBL conferred cytokine hypersensitivity to both GM-CSF (**Figure 6a**) and IL-3 (**Figure 6b**) compared to WT CBL in cell viability assays.^20–23^ In contrast, RE CBL reduced sensitivity to GM-CSF (**Figure 6a**), significantly increasing the GM-CSF EC50 concentration (**Supplementary Figure 6a**) compared to cells expressing WT CBL. Similarly, proliferation of RE CBL expressing cells grown in either GM-CSF or IL-3 was significantly reduced compared to WT or Y371H CBL (**Supplementary Figure 7**). These findings indicate that in addition to EGFR signalling, RE CBL expression also attenuates cytokine receptor signalling more efficiently than WT CBL.

**Figure 6:**
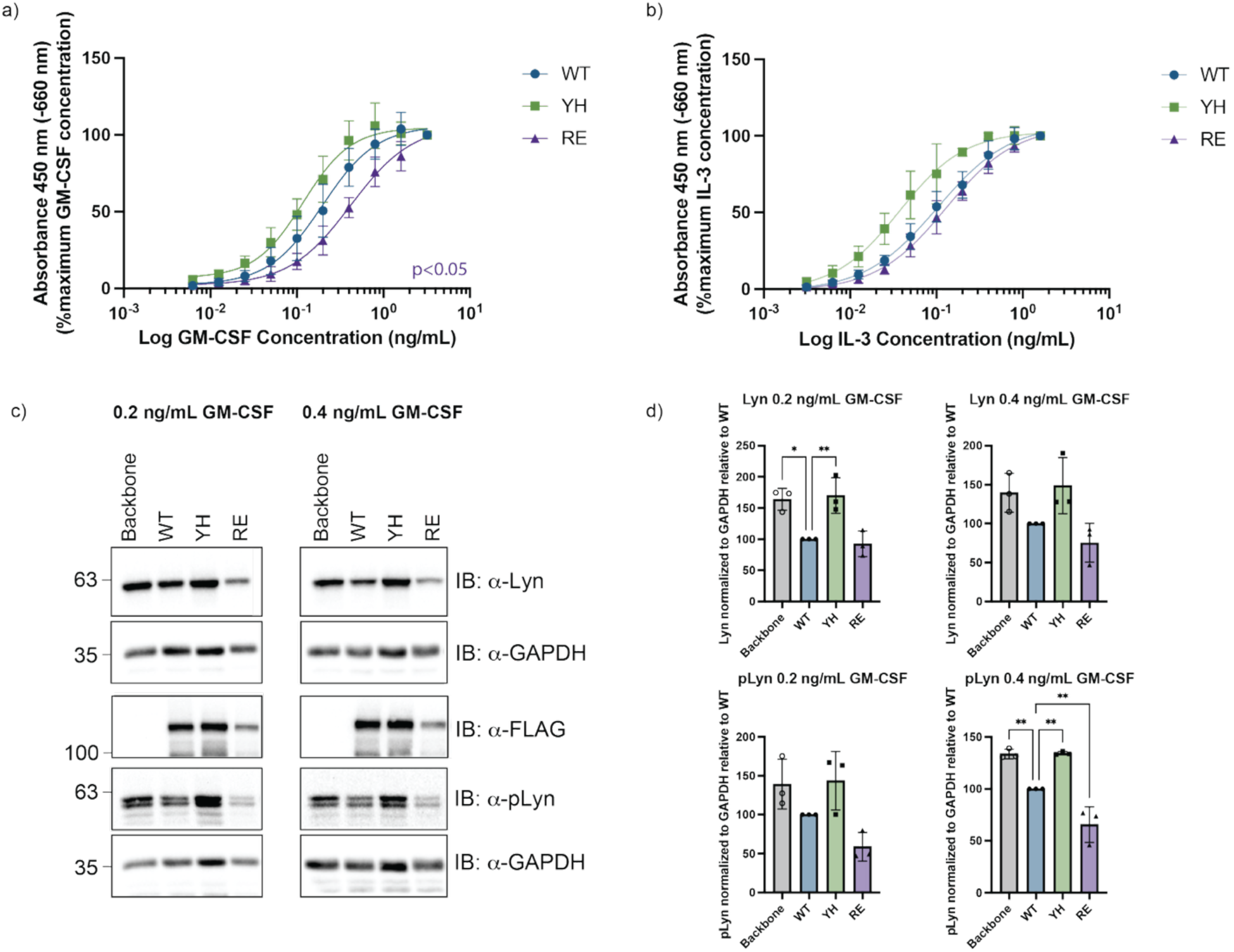
RE CBL reduces cytokine sensitivity of 32D cells and reduces Lyn activation. *Cbl* null 32D cells with induced expression of FLAG tagged WT, RE or YH CBL mutants were grown in increasing concentrations of **a)** GM-CSF or **b)** IL-3 for 72 hours. Cell viability was measured using the CyQUANT XTT assay. The mean absorbance of triplicate wells from four (GM-CSF) or three (IL-3) independent experiments, normalized to the maximum cytokine concentration per line are plotted. Significance levels determined by EC50 comparison (see **Supplementary** Figure 6) **c)** 32D cells were induced to express FLAG tagged CBL and CBL mutants with doxycycline and grown in 0.2 or 0.4 ng/mL GM-CSF for 72 hours. Total lysates were immunoblotted with anti-FLAG (CBL), anti-Lyn and anti-phospho Lyn (pTyr397) and anti-GAPDH as a loading control. **d)** Quantification of Lyn and phosph-Lyn normalized to GAPDH and shown relative to WT CBL. Bars represent the mean normalized values, with individual points showing the values for the independent replicates. Error bars represent the standard deviations between replicates. Comparisons were made to WT CBL by ordinary one-way ANOVA analysis followed by Dunnett correction for multiple comparisons (* p<0.05, ** p<0.002).

It has previously been reported that expression of Y371H CBL promotes greater activation of the CBL substrate, Lyn, in response to cytokine signalling.^20,21^ We examined Lyn and activated Lyn levels in lysates from 32D cells treated with GM-CSF. While YH CBL expressing cells had greater Lyn and pLyn levels compared to WT CBL expressing cells, RE CBL expression reduced active pLyn levels in the presence of both 0.2 and 0.4 ng/mL GM-CSF (**Figure 6c**). A trend towards reduced total Lyn with RE CBL expression in GM-CSF treated cells was also observed (**Figure 6d**). These results indicate that the reduced cytokine responses observed in RE CBL expressing cells is likely due at least in part to reduction in cytokine receptor signalling through activated Lyn.

### Identification of CBL activating compounds

Characterization of the RE CBL protein suggested that small molecules could similarly mimic SLAP2 binding and promote CBL ubiquitin ligase activity to downregulate both receptor and cytoplasmic TK signalling. A high-throughput assay based on an *in vitro* autoubiquitination reaction to measure CBL E3 ligase activity was developed to screen for CBL activating compounds. The assay utilized the *in vitro* “off” state of purified unphosphorylated CBL to identify activators using a 384 well ELISA-based system. In this assay, poly-ubiquitinated proteins were captured by Tandem Ubiquitin Binding Entity (TUBE) coated plates, and relative level of ubiquitination was detected by a luminescent signal.^48^ To scale the use of the assay for the small molecule screen, the time scale for luminescent detection, reagent stability, effects of DMSO and overall work-flow were tested and optimized (**Supplementary Figure 8**).

In the final optimized assay, purified CBL2-436 (including the TKBD-LHR-RING protein regions) was pre-incubated with library compounds for 30 minutes and a reaction master mix containing ubiquitin, ATP, E1 (UBA1) and E2 (UBE2D2) enzymes was subsequently added to allow the ubiquitination reaction to proceed. Following a 90 minute incubation and plate washing, a mixture of a biotinylated polyubiquitin detection reagent and ultrasensitive streptavidin peroxidase polymer was added to detect ubiquitinated species captured by the plate coating. Enhanced chemiluminescent substrate (ECL) was added and the luminescent signal was detected on a plate reader.^164^ Each screen plate included two columns of CBL with pSLAP2 as a positive control, two columns of the negative control CBL alone, and a maximum of 320 compounds assayed per plate (**Figure 7a**). Primary screen hits were determined on a plate-by-plate basis with a cut-off signal of four standard deviations of the compound well mean signal (**Figure 7b**).

**Figure 7:**
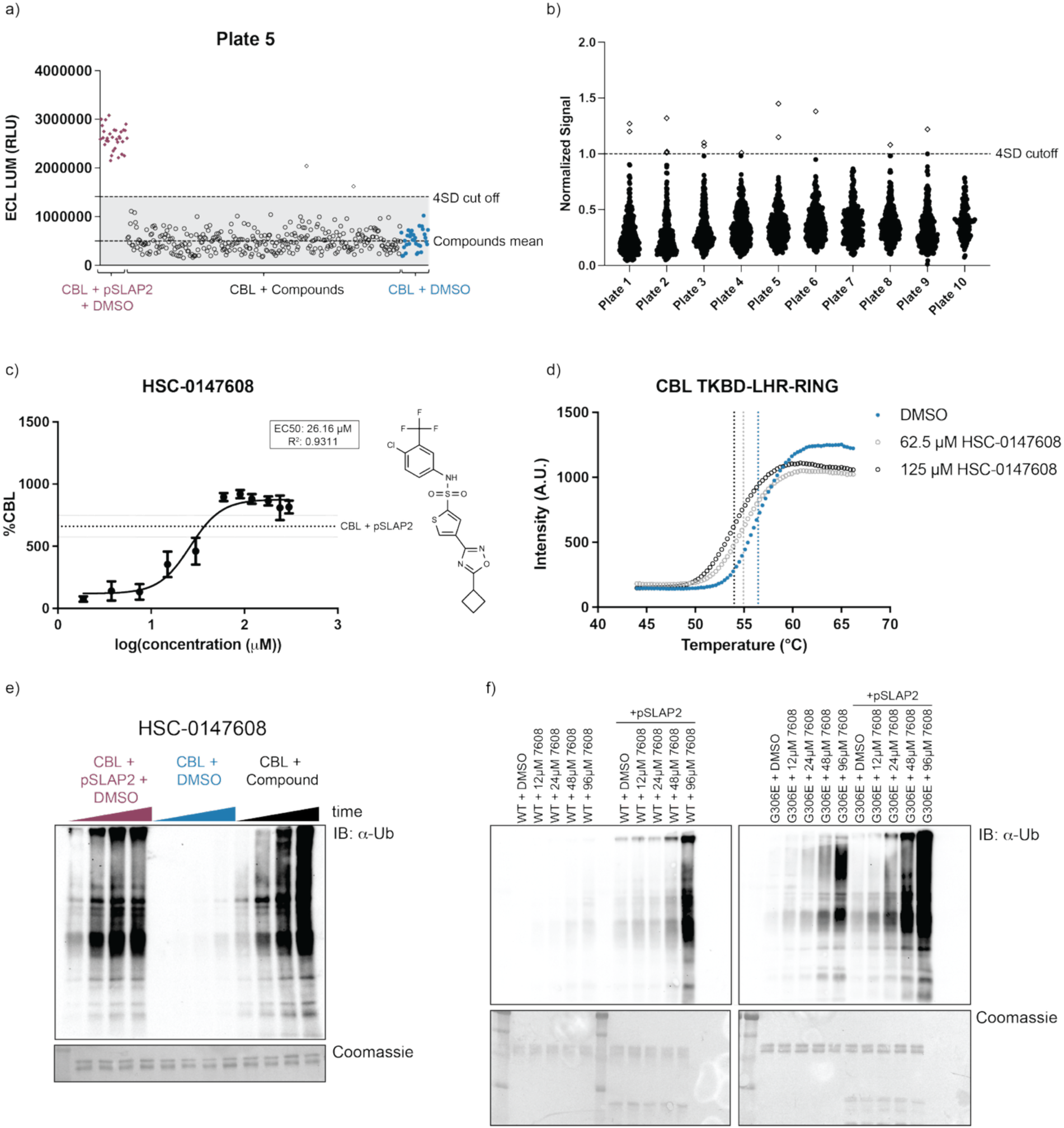
Small molecule screen identifies CBL activating peptidomimetic compounds. **a)** Plot of data from a representative plate with control CBL with pSLAP2 wells shown in red diamonds (left), compounds with open circle points, and negative control of CBL with DMSO (right) as blue circles. Dotted lines show the mean of the compound wells on the plate (lower) and the four standard deviation cut off (upper). **b)** Summary of compound signal from the primary screen normalized by plate to the cut off (indicated at 1). Each dot represents the signal from one compound well. Open dots above the line represent hits. **c)** 11-point dose response with HSC-0147608 (at concentrations range from 1.875 to 300 μM) normalized to average signal for the negative control CBL (giving %CBL). Data from three independent experiments are plotted with level of positive control (CBL + pSLAP2) mean and standard deviation indicated by the horizontal lines. Compound structure is displayed on the right. **d)** Thermodenaturation of CBL2-436 over a temperature gradient in the presence of DMSO or indicated concentrations of HSC-0147608. Vertical lines indicate the temperature of aggregation for DMSO (10%; 56.45°C), 62.5 μM (54.93°C), 125 μM (53.98°C) HSC-0147608. **e)** *In vitro* ubiquitination reactions containing 1200 pmol (48 μM) of HSC-0147608 compound incubated with purified CBL over a time course from 30 to 120 minutes. Reactions were analyzed by immunoblot with anti-ubiquitin. Control reactions with CBL and pSLAP2, or negative control with CBL and DMSO were included. Coomassie stained membranes shows input CBL protein (lower). **f**) Anti-ubiquitin immunoblot of in *vitro* ubiquitination reactions using wildtype (left membrane) or G306E (right membrane) CBL proteins and increasing concentrations of HSC-0147608 or with added pSLAP2 as indicated. Representative immunoblots from two independent experiments are shown.

From the 3000 peptidomimetic small molecules screened, 14 compounds were identified as hits (**Figure 7b**; see **Supplementary Table 1** for screen summary statistics). Ten hits were confirmed in triplicate wells of a single plate in two separate experiments (**Supplementary Figure 9**) and subsequently assayed in triplicate dose response experiments. Four compounds showed some degree of dose dependency, with HSC-0147608 having the most robust response (**Figure 7c** and **Supplementary Figure 10a,b**). In differential static light scattering experiments over a temperature gradient, CBL TKBD-LHR-RING was destabilized in the presence of HSC-0147608 (**Figure 7d**), but not an unrelated compound (**Supplementary Figure 11**) suggesting that HSC-0147608 binds directly to CBL and disrupts autoinhibited conformation. The activity of HSC-0147608 was further validated in a secondary assay in which ubiquitinated species were detected by immunoblot. HSC-0147608 displayed both time (**Figure 7e**) and dose (**Figure 7f**) dependent activation of CBL *in vitro*. In addition, HSC-0147608 activation of CBL was not dependent on the substrate binding site of the CBL TKBD (**Figure 7f**), and was found to have an additive effect in the presence of pSLAP2 (**Figure 7f**). The other confirmed hits had weak effects in immunoblots (**Supplementary Figure 10c**), though this was likely influenced by low solubility. Two of these compounds had high degrees of structural relatedness to HSC-0147608 (**Supplementary Figure 12a**), with low frequency of features in the library overall (**Supplementary Figure 12b**).

## Discussion

CBL is a key regulator of tyrosine kinase signalling pathways that is itself regulated by tyrosine phosphorylation, existing in an autoinhibited state until recruitment to an activated TK, where it is phosphorylated and undergoes a conformational shift into its active form.^11^ SLAP2 binding CBL is also thought to disrupt CBL autoinhibition and promote CBL E3 ligase activity.^48^ In this study, we determined the functional effects of CBL activation via this mechanism using a mutant form of CBL with substitution of A223R and S226E in the CBL-SLAP2 interface of the CBL TKBD, termed RE CBL. RE CBL has enhanced ubiquitin ligase activity both *in vitro* and in cells. Despite mutation in the TKBD, RE CBL interaction with TKs is intact and this mutation does not cause transformation of NIH3T3 cells. Furthermore, we used proximity dependent biotinylation with mass spectrometry (PDB-MS) with MiniTurboID, followed by cellular and biochemical analysis to demonstrate that the RE CBL mutation does not disrupt the CBL interactome, but instead enhances interaction with the CBL substrate, EGFR, as well as early endocytic adaptors. Follow up studies supported that the more efficient activity of RE CBL facilitates greater downregulation of both receptor and non-receptor TKs. Based on these observations we conducted a high-throughput screen and identified a group of structurally related small molecules that mimic SLAP2 binding or the RE mutation by activating autoinhibited CBL, which may have future applications to modulate overactive TK signalling and dysregulated CBL activity in cancer.

We examined the CBL interactome without and with EGF stimulation to understand the effect of the RE mutation in basal and active signalling environments. Within the current literature, the CBL interactome has not been examined through PDB-MS, though has been characterized by AP-MS under varying conditions.^21,25,52,68,69,81–83^ The proximity interactome of RE CBL strongly overlapped with that of WT CBL, including several known CBL interactors, supporting the functional integrity of the RE CBL mutant. The abundance of many proximity interactors was lower with RE CBL compared to WT, likely due to the lower overall MiniTurbo-RE CBL bait levels, particularly in the EGF treated condition. The reduced MiniTurbo-RE CBL bait level is in line with observations that CBL activation results in autoubiquitination and degradation following receptor activation.^84,85^ This also coincides with observations that SLAP2 overexpression causes reduction in CBL levels, supporting the parallel mechanism of RE CBL and SLAP2-mediated CBL activation.^45^

Despite lower RE CBL levels, changes in select proteins in the identified interactomes suggest that the presence of RE CBL shifted the signal transduction and endocytic dynamics of EGFR signalling. For example, RE CBL interacted more readily with EGFR compared to WT CBL across multiple EGF concentrations. Unlike oncogenic mutant CBL proteins, which have both enhanced substrate engagement and sustained signalling,^19,21,25^ RE CBL expression caused enhanced EGFR internalization and attenuation of MEK activation over time. Furthermore, differences in endocytic proteins between WT and RE CBL interactomes, as well as increased modification of EPS15 in RE CBL expressing cells, are consistent with more efficient EGFR ubiquitination, and the subsequent recruitment of endocytic adaptors involved in EGFR downregulation.^79^

We observed more distinct effects of RE and WT CBL expression at lower (5 ng/mL) compared to higher (50 ng/mL) EGF concentrations. This may be due to the predicted open conformation of RE CBL that allows it to overcome an EGFR activation threshold required for stable interaction with the receptor. CBL interacts with the EGFR both directly through TKBD binding to pY1045 of the receptor and indirectly through GRB2 which binds EGFR at pY1068/1086, and both binding methods are required for stable CBL engagement.^39,86,87^ Our findings that RE CBL more readily engaged the receptor at low ligand concentrations, when it is less likely that both EGFR sites are phosphorylated, suggests the active state of RE CBL may bypass the threshold requirement. At higher ligand concentrations differences are likely less discernable as the threshold is passed and both CBL forms are rapidly engaged and activated.

In complementary experiments, we observed that RE CBL led to more efficient downregulation of GM-CSFR signalling, at least in part through reduction of the non-receptor tyrosine kinase, Lyn. Hyperactivation of Lyn downstream of GM-CSFR has been found to be involved in PI3K activation and promote cellular proliferation downstream of oncogenic CBL mutants.^19–22^ These findings suggest that CBL activation through the mechanism elicited by the RE mutation can promote substrate downregulation in multiple clinically relevant pathways.^7,33^ In addition, these data provide insight into the impact of RE CBL expression over extended growth times, and suggest that the reduction in CBL substrate level is sustained.

Previously, biophysical characterization of Tyr371 CBL oncogenic mutants revealed that they are not simply “stuck” in an autoinhibited conformation but rather, these mutations disrupt the native conformation of CBL such that the RING domain is more accessible to E2 binding, and the linker region becomes flexible and does not support stabilization of the active conformation.^88^ Thus, while Tyr371 oncogenic mutant proteins are still able to engage receptors and propagate oncogenic signalling through CBL adaptor function, they do not efficiently ubiquitinate CBL bound substrates.^19,21,23,89^ In contrast, the RE CBL mutation in the TKBD-LHR interface retains Tyr371 and has enhanced phosphorylation, substrate ubiquitination capacity, both indicative of a stabilized active state. The observations that RE CBL promotes both activation in the absence of CBL phosphorylation, and can also support the stabilized activation state by Tyr371 phosphorylation, are in line with the known role of the SLAP adaptor binding to CBL. For example, during T-cell development, SLAP is required alongside CBL for constitutive ubiquitination of the TCRσ chain in the absence of TCR stimulation, and is also important for tyrosine phosphorylation and activation of CBL downstream of activated T-cell and B-cell antigen receptors, RTKs and cytokine receptors.^41,90,91^

With the potential to modulate multiple tyrosine kinase mediated signalling events, we optimized a small molecule screen and identified tool compounds that promote *in vitro* CBL activity. Since an ⍺-helix of the SLAP2 carboxy-terminal tail was found to bind CBL TKBD in a similar region to the ⍺-helix of the LHR in the autoinhibited conformation of CBL,^48^ we reasoned that a peptide-like molecule may be able to similarly bind CBL in this region to promote its activation. We screened compounds from the ChemDiv library that were selected to have structural motifs typical for non-peptide peptidomimetics. We ultimately identified four compounds with dose-dependent CBL activation, with HSC-0147608 showing the most robust and consistent response. Poor solubility profiles of all four compounds prohibited in depth biophysical characterization of their interaction with CBL. However, we were able to demonstrate that HSC-0147608 destabilizes CBL in thermodenaturation experiments, indicating that it is acting directly on CBL. We further predict that in agreement with a destabilizing effect, HSC-0147608 likely disrupts the autoinhibited conformation of CBL. Structural relatedness of three of the four compounds identified with amenability for chemical modification supports viability of future screening of additional related molecules optimized for solubility and drug-like properties and their further characterization of biophysical interactions with CBL.

Promoting CBL activation by the action of the identified compounds and further derivatives may have beneficial effects in several scenarios related to dysregulated CBL activity in cancer. One question is whether these compounds could restore E3 ligase activity of leukemia associated CBL mutant proteins where the RING domain remains intact, but E3 ligase activity is dysregulated. Since the Tyr371 oncogenic mutants destabilize CBL and may retain non-productive ubiquitination activity that contributes to oncogenic signalling, rather than simply “turning off” its E3 activity,^88,92^ it may be warranted to probe whether identified compounds could stabilize the active conformation of Tyr371 mutant CBL. Further, enhancement of CBL autoubiquitination and degradation would also be beneficial to alleviate downstream signalling observed via the maintained adaptor functions of mutant CBL. Indeed, disrupting mutant CBL interaction with TK by “locking” it in a closed confirmation to disrupt association with receptors has recently been reported as a successful strategy in preclinical experiments.^89^

In addition to targeting mutant CBL, promoting wildtype CBL activity may have therapeutic benefits in other contexts, such as in cancers driven by overactive TK signalling. Treatment with CBL activating compounds could also be beneficial in cases with increased CBL Tyr371 dephosphorylation, such as in leukemia where PRL2-mediated CBL dephosphorylation enhances FLT3 and c-Kit signalling. CBL activating compounds may also be therapeutic when CBL is overexpressed, but not phosphorylated, such as was observed in late stage breast cancer.^25,93,94^ There are also several circumstances where CBL appears to be sequestered from overactive TK, particularly EGFR, by other protein-protein interactions.^27–31,35–37^ It would be of interest to determine whether small molecule CBL activators and the presumed change in CBL conformation elicited could disrupt these inhibitory interactions. Finally, combined use of CBL activators and TK inhibitors may provide a strategy to avoid resistance mechanisms. In particular, it would be of interest to investigate potential synergistic effects with dasatinib treatment, which has shown preclinical evidence of attenuating mutant CBL-driven oncogenic signalling.^19–22,95^

In conclusion, detailed characterization of the engineered RE CBL mutant derived from the structural study of the CBL-SLAP2 interaction, revealed a previously unknown mechanism of CBL activation that promotes downregulation of tyrosine kinase signalling, and led to the identification of small molecules that activate CBL E3 function. Our findings provide a foundation for further development of CBL activators, as tools to probe CBL biology as well as therapeutic strategies in preclinical models.

## Acknowledgements

The authors thank Jonathan Sayewich and Dana Wu for experimental support. We thank Drs. David Uehling, Michael Prakesch, and Richard Marcellus from the OICR Drug Discovery Team for library curation and chemical analysis, Dr. Derek Ceccarelli for advice on Uba1 purification, Dr. Brian Raught and Dr. Jonathan St-Germain for BioID/MiniTurboID optimization support, and Dr. Daniela Rotin for the Sleeping Beauty System backbone and transposase plasmids as well as Platinum-E cell line. HeLa Flp-In T-REx cells were kindly provided by Dr. Brian Raught. pDONR221 plasmid was provided by the SPARC Biocentre at SickKids. We also thank the Imaging Facility at SickKids for support with confocal microscopy and the Structural & Biophysical Core Facility at SickKids for support with thermodenaturation experiments. This work was supported by funding from the Cancer Research Society, the Canadian Institutes of Health Research (Project Grant PJT-166034) and the Natural Sciences and Engineering Council of Canada (RGPIN-2019-06485) to CJM. AJT is a recipient of a Canada Graduate Scholarship – Masters, Ontario Graduate Scholarship, and Restracomp Graduate Scholarship (Hospital for Sick Children). SFA and DTH are supported by Cancer Research UK core funding (A29256).

## Author Contributions

AJT performed experiments, acquired and analyzed data, drafted and revised the manuscript. CEM, CDS, DL, and MHE performed experiments, acquired, and interpreted data. LEWG and CF interpreted data. RB, DTH, and SFA provided critical reagents and experimental advice. ACG supervised CEM. CJM supervised AJT, CDS and MHE, analyzed and interpreted data, drafted and revised the manuscript.

## Methods

### Cell maintenance

All cell lines were maintained at 37°C at 5% CO2 in humidified incubators. HEK293T cells were maintained in Dulbecco’s Modified Eagle Medium (DMEM; Wisent) with 10% fetal bovine serum (FBS; Wisent) non-heat inactivated and 1% penicillin/streptomycin. NIH3T3 fibroblast cells were maintained in DMEM with 2.5% calf serum (Wisent). Platinum-E (Plat-E; gift from Dr. Daniela Rotin’s lab) were maintained in DMEM, 10% FBS non-heat inactivated and 1% penicillin/streptomycin, 1 μg/mL puromycin, 10 μg/mL blasticidin S hydrochloride. Parental HeLa Flp-In T-REx cells (gift from Dr. Brian Raught’s lab) were maintained in DMEM, 10% FBS non-heat inactivated, 1% penicillin/streptomycin, 100 μg/mL zeocin and 15 μg/mL blasticidin. After generating expression lines (described below), HeLa Flp-In T-REx cells were maintained in DMEM, 10% tetracycline-free FBS, 1% penicillin/streptomycin, 200 μg/mL hygromycin B and 15 μg/mL blasticidin S hydrochloride. CBL knockout murine myeloblast 32D lines were generated by Dr. Roger Belizaire.^21^ Clone 5A3 was used for the generation of pooled cell lines to re-express CBL and CBL mutants as described below. 32D cells were maintained in RPMI 1640 (Wisent) with 10% heat inactivated tetracycline-free FBS (Wisent) and supplemented with 5 ng/mL IL-3 (Peprotech (Thermo Fisher Scientific) # 213-13) maintained in fresh media every 2-3 days.

### Cell Line Generation

pCDNA5/FRT/TO with a carboxy (C) or amino (N) terminal positioned FLAG-MiniTurboID were cloned through restriction cloning to express wildtype or RE CBL (N terminal into the AscI and NotI sites and C terminal into AfiII and NotI sites). HA-tagged CBL in pCDEF3 (kind gift from Dr. Margo Meyers) was used as a template for WT CBL sequence (the HA tag was not maintained).^47^ The CBL template in pCDEF3 was modified through point mutagenesis using Quik Change II site directed mutagenesis to make the A223R and S226E point mutations (see **Table 1** for primers), and the mutated CBL was subsequently amplified for cloning into the pCDNA5 backbone as for wildtype CBL. For C terminal 3X Flag tagged CBL expression in Flp-IN T-REx lines without the ligase fusion, using gateway cloning, wildtype and RE CBL were introduced into pDEST pCDNA5 vector (obtained from Dr. Anne-Claude Gingras’ lab), after being introduced pDONR221 plasmid (obtained from SPARC Biocentre, SickKids) with attB sites added by PCR from the pCDEF3 template. HeLa Flp-In T-REx cells were co-transfected with pCDNA5 based vectors with the Flp recombinase containing vector, pOG44 (750 ng of each) using Lipofectamine2000 (Invitrogen #11668027) in DMEM with 10% tetracycline-free FBS. Media was changed the next day and two days following transfection, cells were trypsinized and replated in selection media (DMEM with 10% FBS tetracycline-free, 1% penicillin/streptomycin, 15 μg/mL blasticidin S hydrochloride and 200 μg/mL hygromycin B).

**Table 1:**
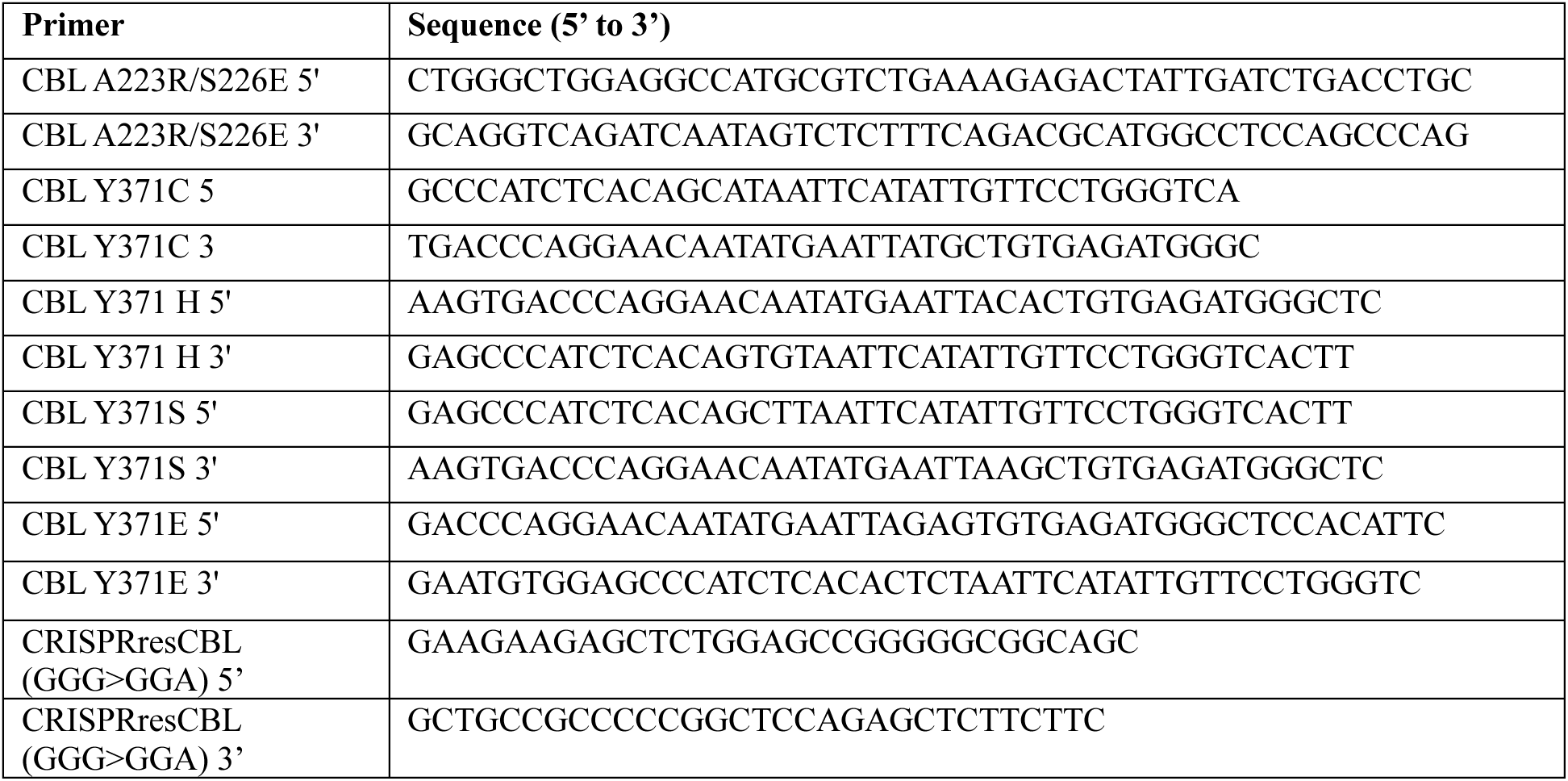
Primers used for site directed mutagenesis.

Following selection, cells were pooled for use in experiments. Control lines were made alongside the CBL fusions using the backbones in the same scheme. pDEST pCDNA5 vectors to express FLAG-tagged CBL were co-transfected in HeLa Flp-In T-REx cells at a 1:10 ratio of expression vector to pOG44 (200 ng of pCDNA5 pDEST vector with 2 μg of pOG44), selected in a similar manner, and clonal cell lines were isolated using cloning rings and screened by immunoblot and immunofluorescence for homogenously expressed CBL. Wildtype CBL CA2, RE CBL CC6 and Control CA4 clones were used in the experiments.

32D lines with CRISPR-mediated *Cbl* knockout were generated by Dr. Roger Belizaire. C terminally 3XFLAG tagged CBL mutants with a point mutagenesis to be resistant to the CRIPSR guide were introduced using the Sleeping Beauty transposase system. CRISPR resistant expression constructs were made via point mutagenesis of pCDEF3 plasmids with wildtype or previously mutagenized CBL to then also change glycine 10 with a silent mutation, GGG>GGA, to disrupt the PAM sequence and avoid targeting by the CRISPR guide (Cbl exon 1 targeting (Cbl KO): FWD: CAC CGT CTT GTC CAC CGT GCA GGG A REV: AAA CTC CCT GCA CGG TGG ACA AGA C) and these were used as a template to clone into modified pSBtet-GFP-Hygromycin.^96^ See **Table 1** for mutagenesis primers. pSBtet-GFP-Hygromycin was modified to introduce AscI and SalI restriction sites, as well as a carboxy terminal 3X FLAG tag, to accommodate that CBL has two internal SfiI sites. CBL expression constructs were introduced into the 32D 5A3 *Cbl* knockout clone through electroporation. 10 million cells were spun at 0.1 rcf for 7 min and resuspended to have 200 μL of cells in RPMI in a 0.4 cm electroporation cuvette (BioRad #165-2088). Cells were mixed with 9.5 μg of pSB expression construct and 0.5 μg of the SB100X vector expressing the Sleeping Beauty transposase (pSBtet-GFP-Hygromycin backbone and SB100X plasmid were a gift from Dr. Daniela Rotin’s lab). Cells were incubated on ice for 5 minutes and then electroporated using a BioRad Gene Pulser Xcell using the exponential protocol at 200V and 950 uF capacitance. Cells were incubated for at least 10 minutes on ice before being transferred to a T25 flask with 10 mL of complete media. Media of cells was changed after approximately 24 hours to select with 600 μg/mL hygromycin B. A control without DNA electroporated was used to assess the selection time. Pooled cell lines were used for experiments. Cells were maintained without hygromycin B and pools were reselected after thaw of new vials every 1-2 months.

### Cell Lysis and Immunoblot

Cells were lysed, cleared by centrifugation, quantified by Pierce BCA Protein assay and prepared for use in pull down experiments or prepared directly for immunoblot. See the **Supplementary Methods** for details on individual experiments. Samples prepared for immunoblot were loaded onto SDS-PAGE gels and resolved prior to transfer to polyvinylidene difluoride membrane. Wet transfer was conducted with 20% methanol. Membranes were blocked in 5% BSA (Roche #10735086001) or 1% fish gelatin (Millipore Sigma #G7765) made up in Tris-buffered saline with 0.05% Tween-20 (TBST) for one hour. Membranes were probed with primary antibody (**Table 2**) overnight at 4°C with gentle rocking. Primary antibodies were diluted in blocking buffer with 0.02% sodium azide. Membranes were subsequently washed with TBST three times for ten minutes each wash. Membranes were incubated in secondary antibody (**Table 2**) diluted in blocking buffer followed by three additional 10 minute TBST washes. Membranes were incubated with prepared enhanced chemiluminescent substrate (ECL) (Revity #ORT2655 and

**Table 2:** Immunoblot antibodies.

| Target | Source | Product code | Dilution |
| --- | --- | --- | --- |
| Anti-mouse IgG, HRP-linked | Cell Signalling Technology | 7076 | 1 in 10 000 |
| Anti-rabbit IgG, HRP-linked | Cell Signalling Technology | 7074 | 1 in 10 000 |
| FLAG M2 | Millipore Sigma | F1804 | 1 in 1000 |
| HA F-7 (His-Ub experiment) | Santa Cruz Biotechnology | sc7392 | 1 in 1000 |
| His6 | Roche | 11 922 416 001 | to 0.5 µg/mL |
| Phospho-Tyrosine 4G10 | Millipore Sigma | 05-321 | 1 in 1000 |
| CBL D4E10 | Cell Signalling Technology | 8447 | 1 in 1000 |
| Streptavidin-peroxidase polymer ultrasensitive | Sigma-Aldrich | S2438-250UG | 1 in 5000 |
| α-Tubulin | Millipore Sigma | T6074 | 1 in 5000 |
| EGFR D38B1 | Cell Signalling Technology | 4267 | 1 in 1000 |
| Transferrin H68.4 | Thermo Fisher Scientific | 13-6800 | 1 in 1000 |
| Erk1/2 | Cell Signalling Technology | 9102 | 1 in 1000 |
| Phospho-Akt (Ser473) | Cell Signalling Technology | 9271 | 1 in 1000 |
| Phospho-MEK1/2 (Ser217/221) | Cell Signalling Technology | 9121 | 1 in 1000 |
| Phospho-Erk1/2 (Thr202/Tyr204) | Invitrogen | 36-8800 | 1 in 1000 |
| Akt | Cell Signalling Technology | 9272 | 1 in 1000 |
| Mek1/2 | Cell Signalling Technology | 9122 | 1 in 1000 |
| pY371 CBL | Kindly provided by Dr. Danny T. Huang's lab |  | 1 in 1000 |
| Ubiquitin P4G7 | BioLegend | 838704 | 1 in 1000 |
| EPS15 D3K8R | Cell Signalling Technology | 12460 | 1 in 1000 |
| HA 16B12 (NIH3T3) | BioLegend | 901501 | 1 in 2000 |
| Lyn C13F9 | Cell Signalling Technology | 2796 | 1 in 1000 |
| Phospho-Lyn (Tyr397)/LCK (Tyr 394)/HCK (Tyr411)/BLK (Tyr 389) (E5L3D) | Cell Signalling Technology | 70926 | 1 in 1000 |
| CBL (32D) | BD bioscience | 610441 | 1 in 1000 |
| GAPDH GA1R | Thermo Fisher Scientific | MA515738 | 1 in 5000 |

**Table 3.**
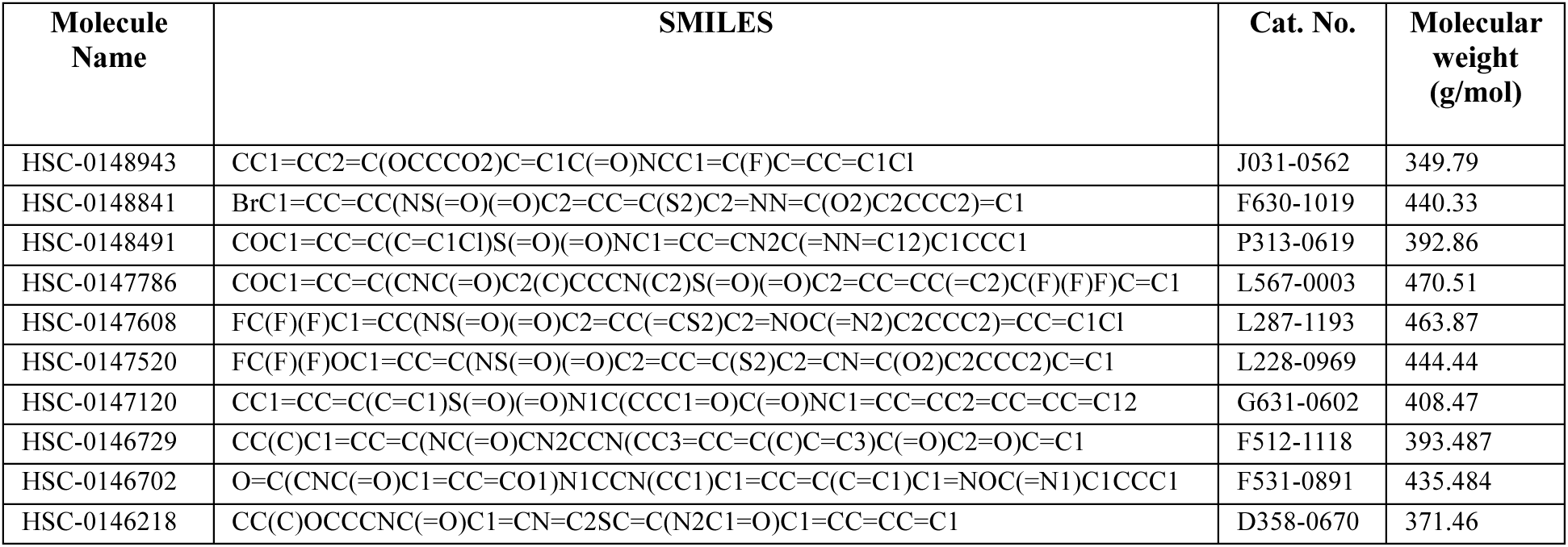
Hit Compound catalogue information.

#ORT2755 prepared with equal parts of these solutions). Membranes probed with Streptavidin-peroxidase polymer ultrasensitive reagent (Sigma #S2438) were blocked in 5% skim milk in TBST and reagent diluted in 5% BSA in TBST. Membranes were incubated with this reagent overnight at 4°C and washed three times with TBST prior to ECL application. Following 2-5 minute ECL incubation, signal was captured on a BioRad Chemi Doc imager (BioRad Laboratories) with multiple chemiluminescent exposures captured, in addition to a colorimetric image of the ladder. Following capture of the chemiluminescent signal, where noted, total protein of lysate membranes was stained with 50% methanol with 0.1% Coomassie Brilliant Blue R-250 dye, and destained in 50% methanol, 10% acetic acid and stained membranes were imaged on the colorimetric setting.

### Immunoblot quantification and data analysis

Band intensities were quantified using the volume tool with background subtraction for similar exposures between replicate blots, with no overexposed bands, using Image Lab (BioRad Laboratories, version 6.0 build 26). Data were normalized as outlined in each experiment. Data was visualized and statistics conducted using GraphPad Prism 10. Where applicable, bars represent the mean normalized values, with individual points showing the values for the independent replicates and error bars represent the standard deviation. Statistical significance was calculated using an ordinary one-way or two-way ANOVA and Dunnett or Tukey’s multiple comparison correction. P value threshold was set as 0.05.

### MiniTurboID proximity dependent biotinylation processing

Stable HeLa Flp-In T-REx cell lines to express carboxy-terminally tagged MiniTurboID-FLAG, wildtype CBL, RE CBL or MiniTurboID control were seeded in DMEM with 1% penicillin/streptomycin and 10% tetracycline-free FBS that was biotin depleted (incubated rocking overnight at 4°C with 1 μL/mL GE-high performance Sepharose Streptavidin-conjugated beads (#17-5113-01)) in 15 cm dishes with 3 plates per cell line per condition. CBL-MiniTurboID expressing cells were induced with 1 μg/mL tetracycline and the control line had tetracycline levels titrated to similar expression levels as the bait with 5 ng/mL tetracycline used for induction, all in fresh media with DMEM with 1% penicillin/streptomycin and 10% tetracycline-free FBS that was biotin depleted. 32 hours after the initial induction, media was changed to DMEM alone with fresh tetracycline to starve the cells for 16 hours. After the starvation period, media was changed to DMEM with 50 μM biotin (Sigma #B4639 made in 30% NH4OH Sigma #221228; and slowly titrated with 12N HCl to make a 20 mM stock solution) for 120 minutes also at 37°C. One set of cells per line (3 plates) were also treated concurrently with 100 ng/mL epidermal growth factor (Millipore Sigma # EA140). After the 120 minute incubation, cells were washed in 30 mL of cold PBS. 800 μL of PBS was then added and cells were gently scraped to harvest (3 tubes per condition). Cells were spun at 500g for 5 min and tubes from the same condition were combined to one pre-weighed 1.5 mL tube with 1 mL of fresh cold PBS. Cells were spun again at 500g for 5 minutes, PBS was aspirated, and pellets were snap frozen on dry ice for at least 5 minutes before being transferred to -80°C. The experiment was conducted in biological triplicate, and all samples were subsequently processed together.

Pellets were first resuspended in a 1:4 ratio of pellet weight to RIPA buffer (50mM Tris-HCL (pH 7.5), 150 mM NaCl, 1% NP-40, 1 mM EGTA, 4.5 mM MgCl2 and 0.4% SDS), supplemented with benzonase (250U/ml, Sigma Aldrich #E1014) and 1X Protease Inhibitor Cocktail (Sigma Aldrich #P8340) and Phosphatase Inhibitor Cocktail 2 (Sigma Aldrich #P5726) and lysed by one freeze-thaw cycle, followed by gentle agitation on a nutator at 4°C for 30 minutes. Cell debris was removed by centrifugation at 16000g for 20 minutes at 4°C. Supernatants were transferred to 1.5 mL microcentrifuge tubes and 25 µL of 60% streptavidin-Sepharose bead slurry (GE Healthcare #17-5113-01) was added. Following overnight incubation at 4°C with rocking, beads were washed with 2% SDS buffer (2% SDS, 50 mM Tris pH 7.5), twice with RIPA buffer and three times with 50 mM ammonium bicarbonate pH 8.0 (ABC).

After removal of the last wash, beads were resuspended in 50 mM ABC (100 µL) with 1 µg of trypsin (Sigma-Aldrich #T6567) and rocked at 37°C for 4h. Then, an additional 1µg of trypsin (in 2µL of 50 mM ABC) was added to each sample and incubated at 37°C with rocking overnight. Beads were centrifuged at 500g for 1 minute, and the supernatant (pooled peptides) were transferred to a new tube. Beads were rinsed with MS-grade H2O (100 µL), and rinse was combined with pooled peptide sample. 10% formic acid was added to the supernatant to final concentration of 2%. The pooled supernatant was centrifuged at 16000g for 5 minutes to pellet remaining beads. 230 µL of the pooled supernatant was transferred to a new 1.5-ml microcentrifuge tube and samples were dried using a vacuum concentrator and stored at -80°C until analysis by mass spectrometry.

Samples were analysed by LC-MS/MS using a Eksigent ekspert™ nanoLC 425 coupled to a TripleTOF™6600 instrument (AB SCIEX, Concord, Ontario, Canada). Nano-spray emitters were generated from fused silica capillary tubing (100 µm internal diameter, 365 µm outer diameter) using a laser puller (Sutter Instrument Co., model P-2000, heat = 280, FIL = 0, VEL = 18, DEL = 2000), followed by packing with C18 reversed-phase material (Reprosil-Pur 120 C18-AQ, 3µm) in methanol using a pressure injection cell. Prior to injection, tryptic peptides were resuspended in 10 µL of 5% formic acid and 2.5 µL (1/4) was further diluted with 3.5 µL of 5% formic acid. 5uL of this volume was loaded onto the LC column using the autosampler and a linear gradient was then delivered at 400nl/min from 2% acetonitrile with 0.1% formic acid to 35% acetonitrile with 0.1% formic acid over 90m. This was followed by a 15m wash with 80% acetonitrile with 0.1% formic acid and equilibration for another 15m to 2% acetonitrile with 0.1% formic acid, resulting in a total of 120m for each run. Samples were analyzed by data independent acquisition (DIA) tandem MS. DIA cycle time was 3.5 seconds, consisting of a 250 ms MS1 scan (400-1250 Da mass range) and 34 x 25 Da SWATH windows (400-1250 m/z mass range). Acquisition of each sample was separated by two wash cycles (using 5% formic acid blanks), each with 3 rapid gradient cycles at 1500 nl/min over 30 min, and followed by a 30 min mass calibration run (using BSA).

### Mass Spectrometry Analysis

Raw files were stored using the ProHits laboratory information management system (LIMS) platform (2) and analyzed using the directDIA module in Spectronaut (18.1) to identify and quantify protein group intensities. The settings were as follows: Trypsin/P as specific enzyme; peptide length from 7 to 52; max missed cleavages 2; toggle N-terminal M turned on; Carbamidomethyl on C as fixed modification; Oxidation on M and Acetyl at protein N-terminus as variable modifications; FDRs at PSM, peptide and protein level all set to 0.01; minimum fragment relative intensity 1%; 3–6 fragments kept for each precursor, normalization and imputation off.

SAINTq scoring of Spectronaut-calculated intensities identified the probability of true interactors from the background contaminants using 6 controls of the MiniTurboID-3X FLAG (3 without EGF treatment and 3 with EGF treatment from 3 biological triplicate experiments run alongside the CBL-MiniTurboID expressing cells).^97,98^ Interactors were scored as significant based on a BFDR cutoff of 0.01 and below. Note that interactors are listed by gene name.

Data was visualized using ProHits-viz to generate dotplots.^99^ Interactors were searched for known interactions with CBL using BioGRID (2024/01/23) and through PubMed searches for “CBL AND *prey*”. Categorizations of interactors were made using GeneCards and Uniprot analysis in addition to gene ontology terms and other characterizations identified using g:Profiler. Further analysis of subsets of data were also conducted using g:Profiler and STRING (V 12.0).^100,101^ To compare interactors by EGF condition or WT versus RE, all interactors that met the BFDR cutoff of 0.01 and below for either condition being compared were considered. A Student’s T test comparison between the three replicates was made for each interactor in Excel (2 tail, homoscedastic – assuming equal variance). The ratio of the average spectral intensity across the three replicates for each interactor was taken comparing that for RE CBL to WT or EGF to no EGF. Log2 of that ratio was taken and plotted against the -log10 of the p value from the T test. Statistics were conducted in Excel. Scatter plots were generated using Python 3 using plotly.

### NIH3T3 Transformation assay

HA-tagged CBL and CBL mutants were cloned into the pBABE-puro retroviral expression vector using restriction cloning with BamHI and SalI. Wildtype, Y371mut and A223R/S226E CBL (see **Table 1** for mutagenesis primers) were amplified from pCDEF3, and 70Z CBL was amplified from pCDNA3.1 to add restriction sites and maintain the HA tag. For retroviral production, Plat-E cells were plated in DMEM with 10% FBS. The next day, cells were transfected with 9 μg of pBABE-puro expression constructs using Lipofectamine® LTX with Plus reagent (Thermo Fisher #15338030) per the manufacture’s protocol. The day after the Plat-E cells were transfected, media was changed and (DMEM with 10% FBS) and 300 000 NIH 3T3 cells were seeded in 10 cm dishes with one dish per condition (CBL mutant) in DMEM with 2.5% CS. 48 hours after the Plat-E transfection, the cell supernatant was harvested to collect the retrovirus and filtered using 0.45 μm PVDF syringe filter (Millipore Sigma #SLHVR33R5). 8 mL of viral supernatant and 8 mL of media (in DMEM with 2.5% CS) were added to the NIH3T3 cells with polybrene (Millipore Sigma #TR-1003-G) to a final concentration of 6 μg/mL per dish. Fresh media was added to the Plat-E cells (DMEM with 10% FBS) and the infection was repeated the next day at 72 hours post-transfection. The day after the second retroviral transduction, the media of the NIH3T3 cells was changed to DMEM with 2.5% CS and 2 μg/mL puromycin for 3 days with media changes each day. Following the third day of selection, the infected cells were passaged and 10000 to 25000 were seeded in at least 4 wells of 6 well plates in DMEM with 2.5% CS with 1% penicillin/streptomycin. Media was changed approximately every 3 days. At day 9 following the cell seeding, one set of replicates was lysed. Cells were washed in cold PBS lysed in 50 μL of lysis buffer (50 mM Tris pH 7.5, 150 mM NaCl, 1% Nonidet P-40, 10% glycerol, supplemented with a mini cOmplete EDTA-free protease inhibitor tablet (Roche #11836170001)). Cells scraped into lysis buffer and lysed on ice for 30 minutes before clearing through 20817 rcf centrifugation for 15 minutes at 4°C. 30 µg of lysate was prepared with 2X sample buffer (125 mM Tris pH 6.8, 4% SDS, 20% glycerol, 0.715 M β-mercaptoethanol, + bromophenol blue) for immunoblot. 13 days following the seeding of the 6 well plates, the colonies were stained. Media was aspirated from cells and plates were washed twice with PBS (with Ca^2+^/Mg^2+^). Cells were fixed in methanol on ice for 10 minutes. 1 mM sulforhodamine B sodium salt (Sigma Aldrich #S1402-16) solution (prepared in 1% acetone) was added to the plates, which were subsequently incubated for 30 minutes on an orbital shaking platform wrapped in foil. Plates were washed in MilliQ water three times to remove excess dye and left to dry overnight. Stained plates were imaged on a Zeiss AxioZoom microscope on the tile setting with 100% brightness and 7.8 ms exposure, with tiled images taken across the full 6 well plate (X = 130000 μm and Y = 80000 μm). Images were fused with minimal overlap of 1% and maximal shift of 1% on the Zeiss software and exported as tiffs. Brightness and contrast adjustments were identically applied to representative images shown in Adobe Photoshop 2024 (-26 brightness, 75 contrast). After imaging, dye was extracted in 2 mL of 10 mM Tris pH 7.5 by shaking for 30 minutes. 200 μL of the extracted dye from each well was plated in triplicate and the absorbance was measured at 564 nm on a Molecular Devices VersaMax micro plate reader.

For each condition, the triplicate well readings were averaged. The mean of those averages for the technical triplicate wells of the 6 well plates were plotted. The experiment was conducted with biological duplicates in this manner with an individual replicate shown.

### XTT viability assay

CyQUANT XTT (Invitrogen #X12223) assay was used to assess 32D cell viability after 72 hour incubation with varying conditions indicated in figures. 32D cells with *Cbl -/-* and inducible expression of wildtype or mutant CBL were washed twice with PBS with spins at 0.1 rcf for 7 min and resuspended in media (RPMI with 10% tetracycline-free FBS), 500 µL was taken to count by ViCell XR cell counter (Beckman-Coulter) and diluted to 30 000 cells/mL. Media was made up to have 2X doxycycline (for 500 ng/mL final) and 2X concentrations of cytokine (recombinant murine IL-3 PeproTech #213-13; recombinant murine GM-CSF eBioscience #BMS325; made in serial dilution) to have 50 µL of media and 50 µL of cells added per plate with 1500 cells seeded per well. Triplicate wells were seeded for each condition for each cell line, including triplicate blank wells per plate. Cells were incubated for 72 hours at 37°C with 5% CO2. Following the incubation, the XTT assay reagent was prepared per manufacturer’s protocol and 70 µL of prepared reagent was added per well. Plates were incubated at 37°C for 4 hours, spun for 1-5 min at 4000 rpm and read at 450 nm and 660 nm (background). Averages were taken for the triplicate wells of the corrected 450 nm absorbance, with 660 nm values subtracted as well as background from the blank. Biological triplicate experiments were conducted (four experiments for GM-CSF) and data combined from the averages of the triplicate wells of each individual experiment. Data were normalized for each cell line to the highest cytokine concentration and plotted as a %maximum of the normalized values. Concentration values were transformed logarithmically. Nonlinear regression (log(agonist) vs. response --Variable slope (four parameters)) was performed to calculate EC50 values for individual experiments. The EC50 values from the triplicate experiments were compared to wildtype CBL by ordinary one-way ANOVA analysis followed by Dunnett correction for multiple comparisons. Data were analyzed in GraphPad Prism 10.

### Protein expression

Residues 2-436 of CBL were cloned in frame into pGEX4T-1 to express glutathione-S-transferase (GST)-CBL TKBD-LHR-RING. Residues 29-261 of human SLAP2 were cloned in frame into a modified pET32a vector (Wybenga-Groot & McGlade, 2021) to express thioredoxin-tagged (Trx)-His(6)-hSLAP2.^48^ Human Uba1 (ORF: BC013041) was expressed from the pETM-30 backbone (Dr. Frank Sicheri’s lab). See **Supplementary Methods** for purification details.

### Primary screen

The compound library included 3000 compounds curated from the ChemDiv Nonpeptide Peptidomimetic PPI library (stock concentration 1 mM in DMSO) for screening using E3Lite 384 well plates (LifeSensors #UC107). Compounds were used from stock libraries at the SPARC Biocentre screening facility at SickKids. Columns 1-2 for each plate were a positive control of 10 pmol wildtype CBL with 30 pmol of pSLAP2 and DMSO (0.3 or 3% final). Columns 23 and 34 were the negative control of 10 pmol wildtype CBL with DMSO. Compounds were present in columns 3-22 with 30 μM of compound per well (15 μM final in the reactions).

Assay plates were thawed at room temperature prior to the assay setup. CBL solutions (10 pmol in PBS + 5 mM βME) were prepared and kept on ice before being added to a 96 well reservoir plate (Corning 96 V-bottom #3363). Plates were washed 2X with 50 μL of PBS per well using the BioTek Plate washer (75 μL/well at a flow rate of 7). 20 μL of CBL mixture was added to the assay plate using a BRAVO liquid handler with a 96 tip head. Compounds were added using an Echo 550 Acoustic Liquid handler (600 nL dispensed). The plates were spun briefly and then incubated at room temperature for 30 minutes with a lid. Using the Multi drop Combi with a metal tip head, 20 μL of ubiquitination reaction master mix (giving final concentrations in 40 μL of: 1X reaction buffer (20 mM Tris pH 7.5, 1 mM MgCl2, 2 mM DTT), 625 μM ATP (25 nmol/well), 7.61 µM ubiquitin (0.3 nmol/well, R&D Systems #U-100H), 26.25 nM E1 (1.05 pmol/well, UBE1 Life Sensors #UB-0101-0050), 37.5 nM E2 (1.5 pmol/well, UBE2D2 R&D Systems #E2-622-100)) was added to each well. The plates were spun briefly and incubated for 1.5 hours at 30°C shaking with a lid. Using the BioTek plate washer, the plates were then washed 3X with 50 μL of PBS-T (0.1% Tween-20) per well at a flow rate of 7. Using the Muti drop Combi, 25 μL of antibody mixture (1:1000 dilution of a ubiquitin binding domain Detection reagent 1, Life Sensors; 1:10 000 Streptavidin-peroxidase polymer ultrasensitive, Sigma #S2438-250UG) made up in PBS-T with 5% BSA filtered through a 0.22 μm PES SteriCup filter; Millipore Sigma #S2GPU05RE) was added to each well. Plates were spun briefly and incubated for 1 hour at room temperature with agitation. Using the BioTek plate washer, the plates were then washed 3X with 50 μL of PBS-T (0.1% Tween-20) per well. Using the Multi drop combi, 25 μL of ECL solution (Millipore Immobilon Western Chemiluminescent HRP Substrate #WBKLS0500) was added to each well, the plates were briefly spun and incubated for 5 minutes at room temperature before the luminescent signal was read using a Synergy Neo plate reader (Top filter cube: #3 – Single PMT LUM, Bottom filter cube: # 114 – LUM 1536; gain of 120 and optics position = Top). CBL mixture, ubiquitination reaction master mix and ECL were made once per screen day. Two antibody mixture were made each screen day (used for 4-5 plates each; at most 40 minute old solution added). Quality of assay runs during the optimization and throughout the screens were monitored throughout the screen using Z’ score calculations of the control wells (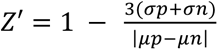, where *σp* and *σn* represent the standard deviation of the luminescent signal from the positive and negative control wells respectively, and *μp* and *μn* represent the means of the signals for the positive and negative control wells respectively).

### Hit classification and validation experiments

Hits were called from the primary screen on a plate-by-plate basis as any compound giving a signal at least 4 standard deviations from the mean of the signal from compound wells on the plate. Two hit validation experiments were run with all hit compounds on one plate, including three wells per hit compound. Hit validation experiments were run as described for the primary screen. Confirmed hits were called based on the percent activation (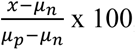, where *x* is the signal from one compound well and *μp* and *μn* represent the means of the signals for the positive and negative control wells respectively), with confirmed hits having a percent activation of 20% or over in at least 2 of 3 wells in at least one of the two replicate experiments.

### Compound dose response

11-point dose response curves were run for the 10 confirmed Peptidomimetic hit compounds (at concentrations of: 1.875, 3.75, 7.5, 15, 30 (screen concentration), 60, 90, 120, 180, 240, 300 μM). The concentrations are based on a 20 μL volume which the compounds were added to, while the final concentrations in the reactions are halved as each reaction ran at 40 μL following the addition of the master mix. This data encompasses three independent experiments, where each concentration had duplicate wells per plate. Each plate also included wells of the negative control, CBL alone, and the positive control, CBL + pSLAP2. The log of the concentrations were plotted on the x-axis. To more accurately combine the three replicates, the relative luminescent values were normalized to the average CBL signal for the respective plates and then multiplied by 100 giving the %CBL signal. The mean of 6 values was plotted with the standard deviations against the log of the concentrations on the y-axis. The horizontal lines indicate the average %CBL for the positive control, CBL + pSLAP2, with the upper and lower lines showing the boundaries of the standard deviations of the controls across replicates. Non-linear regression with variable slope was run using GraphPad Prism 6 to determine the EC50 values. Compounds used in the dose response experiments were purchased fresh from Molport and prepared in DMSO to 10 mM.

### Small scale *in vitro* reactions

E3Lite plate assays in **Figure 1** were conducted as previously described.^48^ For immunoblot experiments, 25 μL reactions were set up in 1.5 mL tubes with 20 pmol CBL (800 nM), 60 pmol pSLAP2 (2400 nM where indicated), 1 mM ATP, 12.18 μM ubiquitin, 42 nM E1 (E1 was either from LifeSensors #UB-0101-0050 or purified in lab), 60 nM UBE2D2, in reaction buffer with 20 mM Tris pH 7.5, 1 mM MgCl2, 2 mM DTT and indicated compound concentrations, with DMSO controls run with the corresponding %DMSO. Reactions were incubated for indicated times at 30°C with 400 rpm shaking in a thermomixer. Ubiquitination reactions were terminated through the addition of 25 μL of 2X sample buffer (125 mM Tris pH 6.8, 4% SDS, 20% glycerol, 0.715 M β-mercaptoethanol, bromophenol blue) and the reactions were boiled for 10 minutes. 25 μL of each reaction mixture was resolved on an SDS-PAGE 12.5% gel and immunoblotting conducted as described above with blocking and antibodies prepared in 5% BSA solutions.

### Thermodenaturation

Using the StarGazer-2, temperature scan experiments were conducted at a temperature range of 20°C to 95°C with a ramping of 1°C per minute. 246 images with 50 ms exposure were taken over 74 minute experiments. Intensity data from the differential light scattering was measured. 0.4 mg/mL CBL2-436 diluted in 500 mM NaCl, HEPES pH 7 was used to assess HSC-0147608 at indicated concentrations or in DMSO (final of 5-10% DMSO in all conditions). The CLK1 inhibitor TG003 (Alfa Aesar #J64690; made to 10 mM stock in DMSO) was used as a negative control. Temperature of aggregation (Tagg) was calculated using the Stargazer-AIR software with a Boltzmann Regression.

