## Supplementary Figures and Methods for "Targeting CBL ubiquitin ligase activation to downregulate tyrosine kinase signalling"

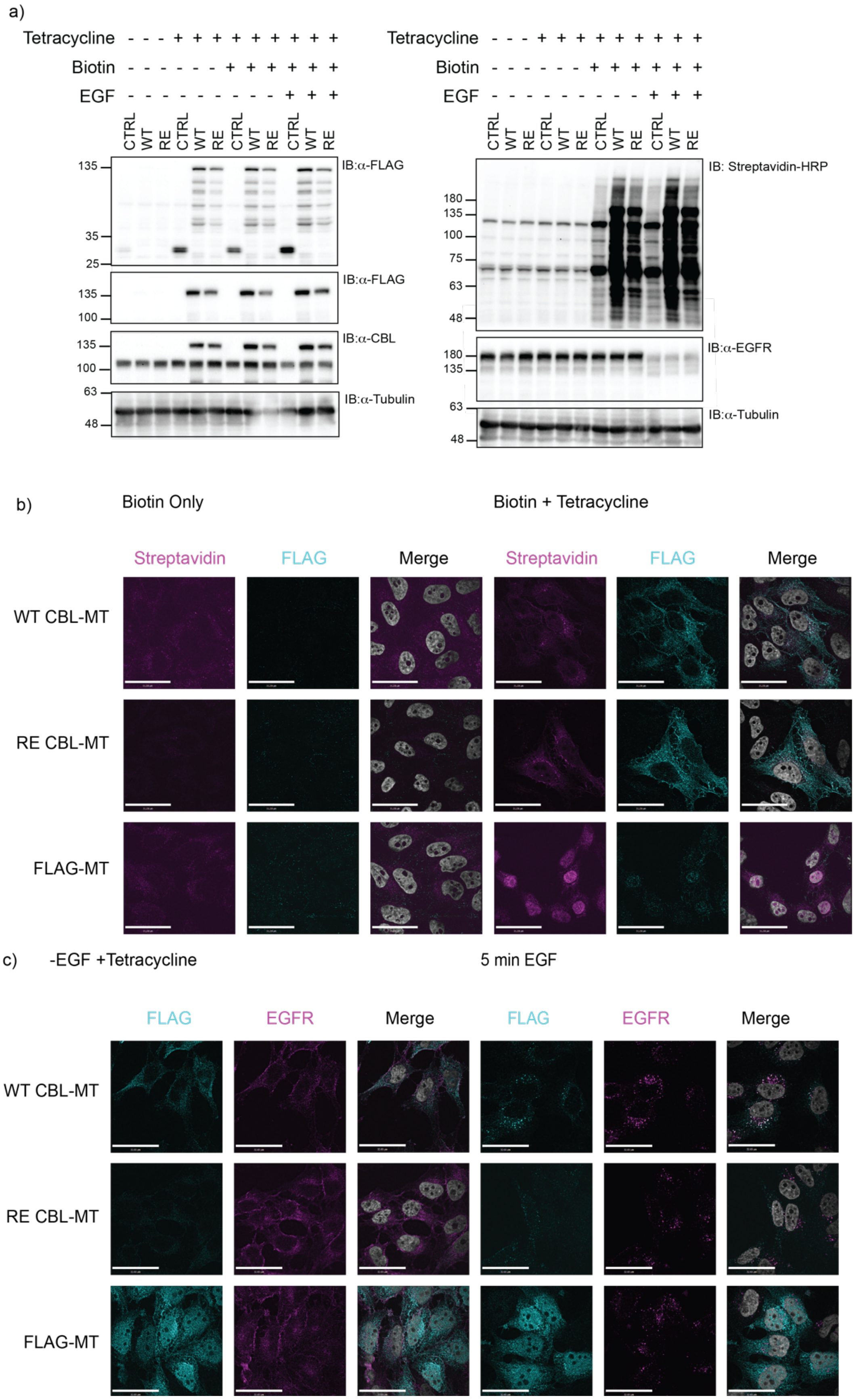

a)
Tetracycline - - - + + + + + + + + +
Biotin - - - - - - + + + + + +
EGF - - - - - - - - - + + +
CTRL WT RE CTRL WT RE CTRL WT RE CTRL WT RE
135
35
25
IB:α-FLAG
135
100
IB:α-FLAG
135
100
IB:α-CBL
63
48
IB:α-Tubulin
Tetracycline - - - + + + + + + + + +
Biotin - - - - - - + + + + + +
EGF - - - - - - - - - + + +
CTRL WT RE CTRL WT RE CTRL WT RE CTRL WT RE
IB: Streptavidin-HRP
180
135
100
75
63
48
IB:α-EGFR
180
135
63
48
IB:α-Tubulin
b)
Biotin Only
Biotin + Tetracycline
Streptavidin
FLAG
Merge
Streptavidin
FLAG
Merge
WT CBL-MT
RE CBL-MT
FLAG-MT
c)
-EGF +Tetracycline
5 min EGF
FLAG
EGFR
Merge
FLAG
EGFR
Merge
WT CBL-MT
RE CBL-MT
FLAG-MT

**Supplementary Figure 1: Validation of the CBL-MiniTurboID cell lines**

**a)** Representative blot of the MiniTurboID workflow conditions for WT or RE CBL with carboxy-terminal MiniTurbo-FLAG tag and the control line expressing MiniTurbo-FLAG. Lysates were collected and resolved by SDS-PAGE and immunoblotted for FLAG (to show the MiniTurbo ligase and tagged CBL proteins), CBL (upper bands show tagged protein and the lower band is the endogenous CBL), tubulin as a control (left panel) probed with streptavidin-HRP and anti-EGFR, or anti-tubulin as a loading control (right panel). **b)** To validate that biotin labelled proteins were localizing near to the carboxy terminally tagged FLAG-MiniTurbo-CBL, the HeLa cells expressing wildtype or RE CBL fused to the MiniTurbo ligase (denoted as MT) were seeded on coverslips and treated as planned for the proteomics experiment and stained by immunofluorescence for FLAG or streptavidin to show the localization of biotinylated preys of the MiniTurbo tagged CBL baits. **c)** The MiniTurbo-CBL were seeded and induced as in **b)**, and then treated without or with 5 minutes of 100 ng/mL EGF. Fixed and permeabilized cells were then co-stained for FLAG and EGFR to show co-localization of the MiniTurbo tagged CBL proteins and EGFR in response to EGF. EGFR staining is shown in magenta and FLAG staining in cyan, with DAPI nuclear staining shown in gray in the merged images.

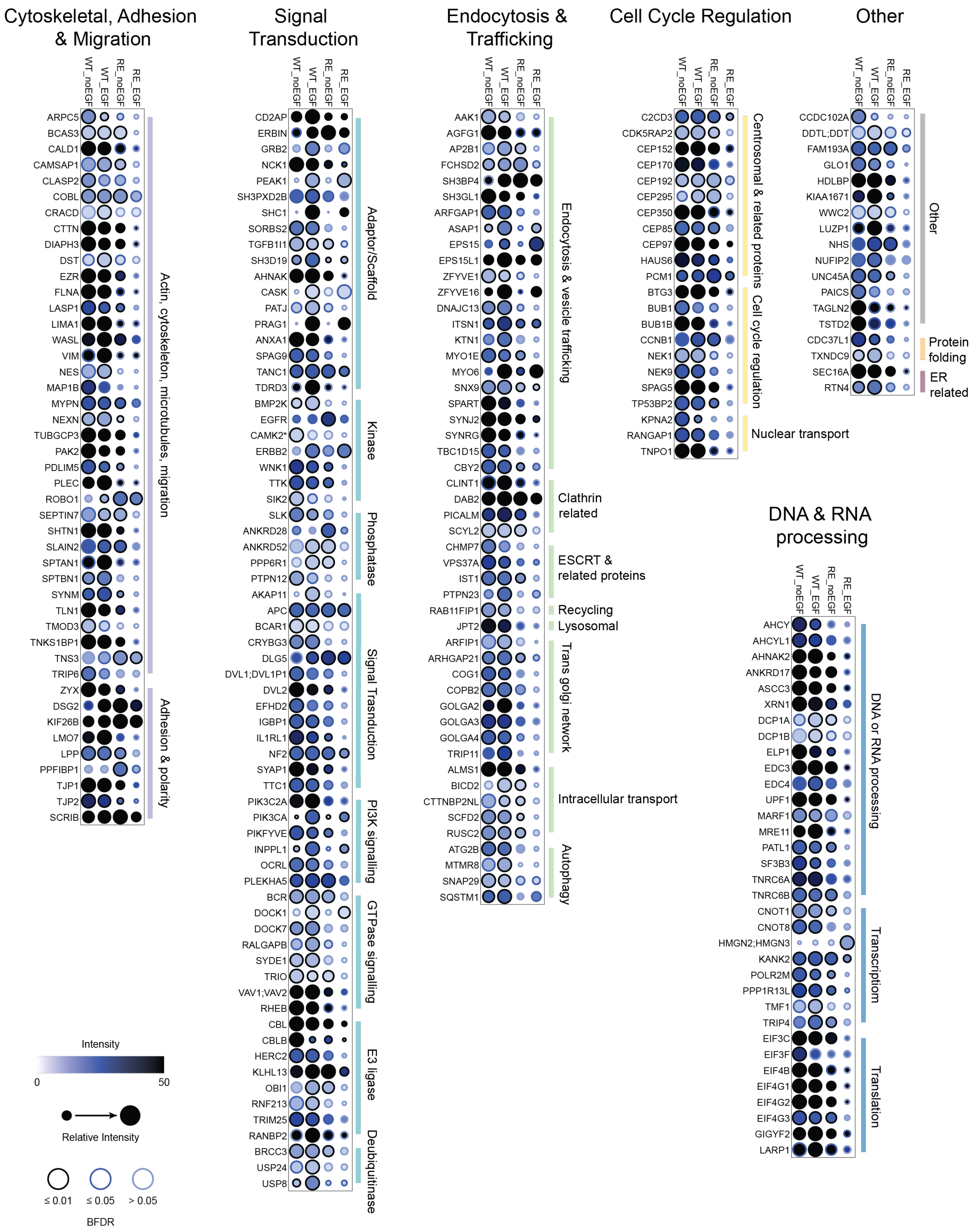


**Supplementary Figure 2: Comparing wildtype and RE CBL proximity interactor abundance across groups of proteins identified**

Overall dot plots of the categorizations identified for the HeLa proximity dependent biotinylation experiments with WT and RE CBL. The different conditions are listed in columns, with each of the rows indicating an identified interactor. Subcategories are listed to the right with groupings denoted by coloured lines. The data was filtered with a BFDR score of 0.01 and a secondary filter of 0.05. As shown in the legend, the circle outlines indicate the significance values. The intensity of the blue is relative the abundance (spectral intensity). The size of the circles indicates the relative abundance of an individual interactor across the four conditions. CAMK2* indicated as nonspecific identification of CAMK2B; CAMK2G; CAMK2D; CAMK2A was noted. Dot plots were generated using ProHits-viz.

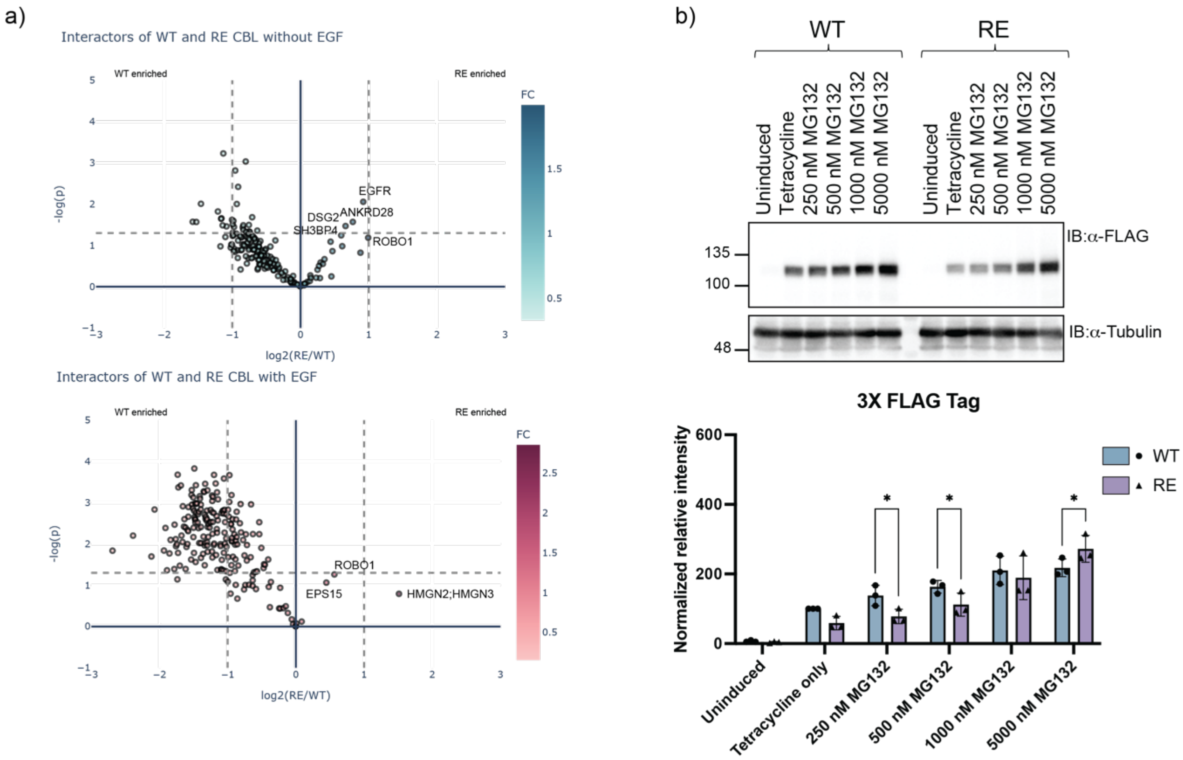


**Supplementary Figure 3: Interactor abundance likely influenced by RE CBL autoubiquitination and degradation**

**a)** To compare wildtype (WT) and RE CBL interactors, preys identified were selected by significance cutoff of a BFDR of 0.01 and below without (top; teal) or with (lower; red) EGF treatment in either the WT or RE CBL interactomes. Interactors that were more abundant with RE CBL are on the right of the graphs, while interactors that were more abundant with WT are found to the left. The triplicate spectral counts of each prey in WT compared to RE CBL were analyzed by T tests with the -log of the p value plotted on the y-axis. The $\log_2$ of the ratio of RE to WT is plotted on the x-axis. Vertical lines indicate a fold change of 2 and over in either direction. The horizontal dotted line indicates a p value of 0.05, points above this line are interactors with a p value scoring less than 0.05. Select interactors are labeled. The colouring of the dots is relative to the ratio of RE to wildtype CBL per the scale on the right. **b)** Protein lysates of HeLa Flp-In T-REx cells expressing C terminally tagged 3XFLAG tagged WT or RE CBL treated with increasing concentrations of MG132 were analyzed as in **Figure 2c**).

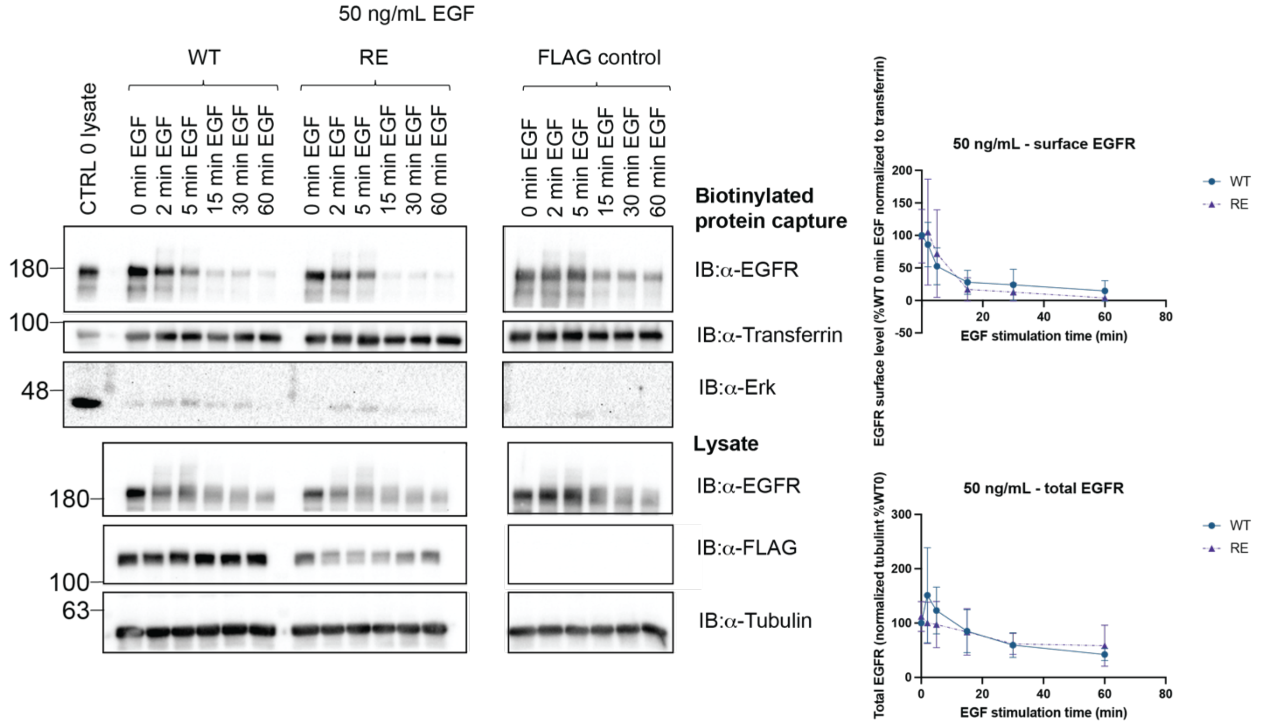


**Supplementary Figure 4: Surface level EGFR levels did not significantly differ when stimulated with high EGF**

Assessment of surfaces level EGFR through surface level biotinylation with HeLa cells induced to express WT or RE CBL for 8 hours in full serum and serum starved for and addition 16 hours. Following induction cells were treated with 50 ng/mL EGF at 37°C for the indicated times. Cell surface proteins were biotinylated and captured from cell lysate, followed by immunoblot of samples. EGFR, transferrin as a positive control for surface proteins, and the intracellular protein Erk as a negative control were assessed for the surface level protein capture. Cell lysates from the same experiments were also run. Figures show representative blots from independent triplicate experiments. In the biotinylated protein capture blots, EGFR and transferrin signals were quantified. In each experiment, EGFR levels were normalized to the transferrin level and then to the signal for WT CBL at 0 ng/mL EGF (for both WT and RE) and the mean and standard deviation from three independent experiments are shown. Two-way ANOVA analysis (with Tukey correction) not a significance difference in EGFR levels with 50 ng/mL EGF treatment. Similarly, lysate blots were quantified for EGFR and tubulin. EGFR levels were normalized to tubulin level and then to the signal for WT CBL at 0 ng/mL EGF (for both WT and RE) and the mean and standard deviation from three independent experiments are shown. Two-way ANOVA analysis revealed no significant difference.

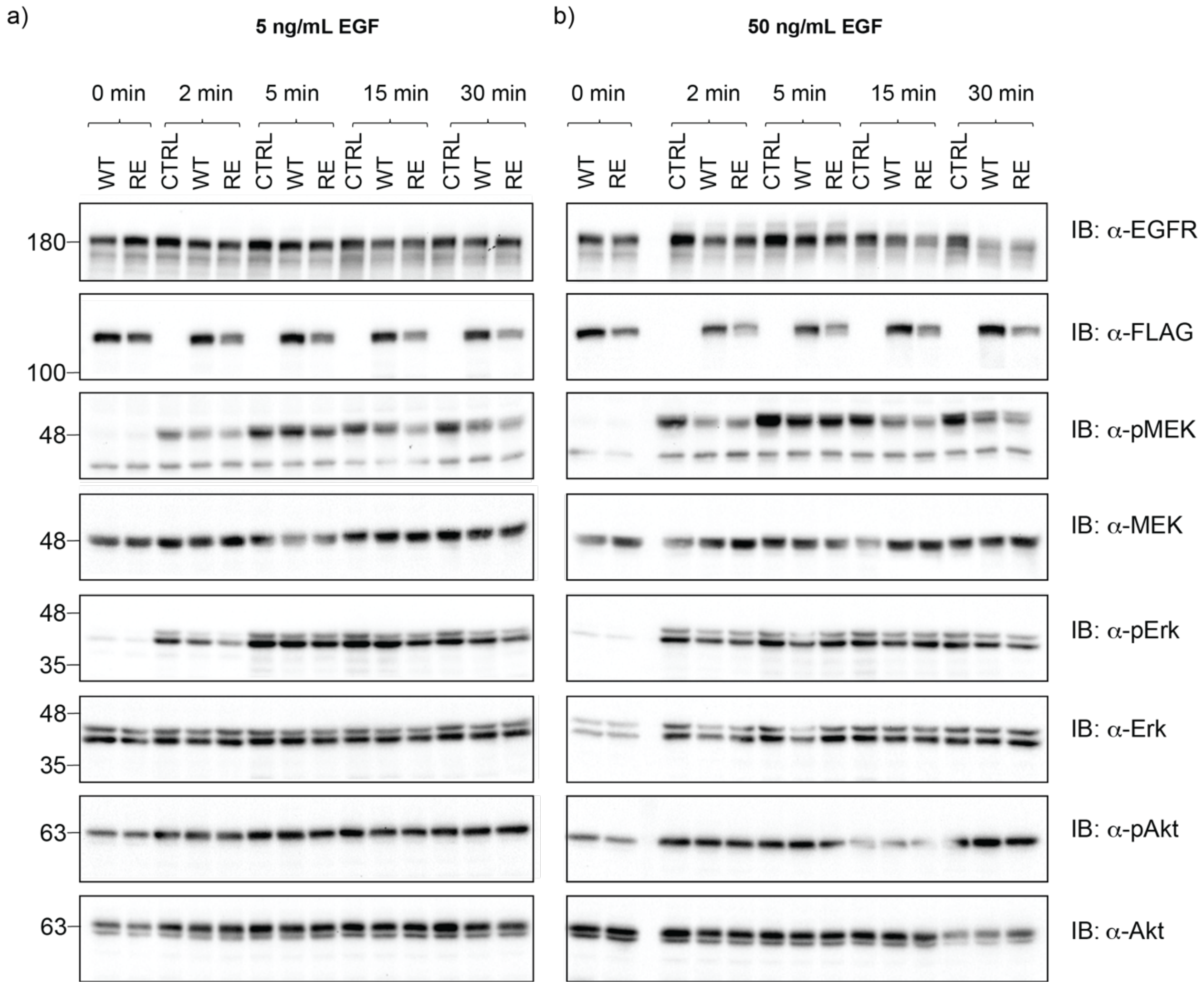
a)
5 ng/mL EGF
b)
50 ng/mL EGF
0 min
2 min
5 min
15 min
30 min
WT
RE
CTRL
180
100
48
35
63
IB: α-EGFR
IB: α-FLAG
IB: α-pMEK
IB: α-MEK
IB: α-pErk
IB: α-Erk
IB: α-pAkt
IB: α-Akt

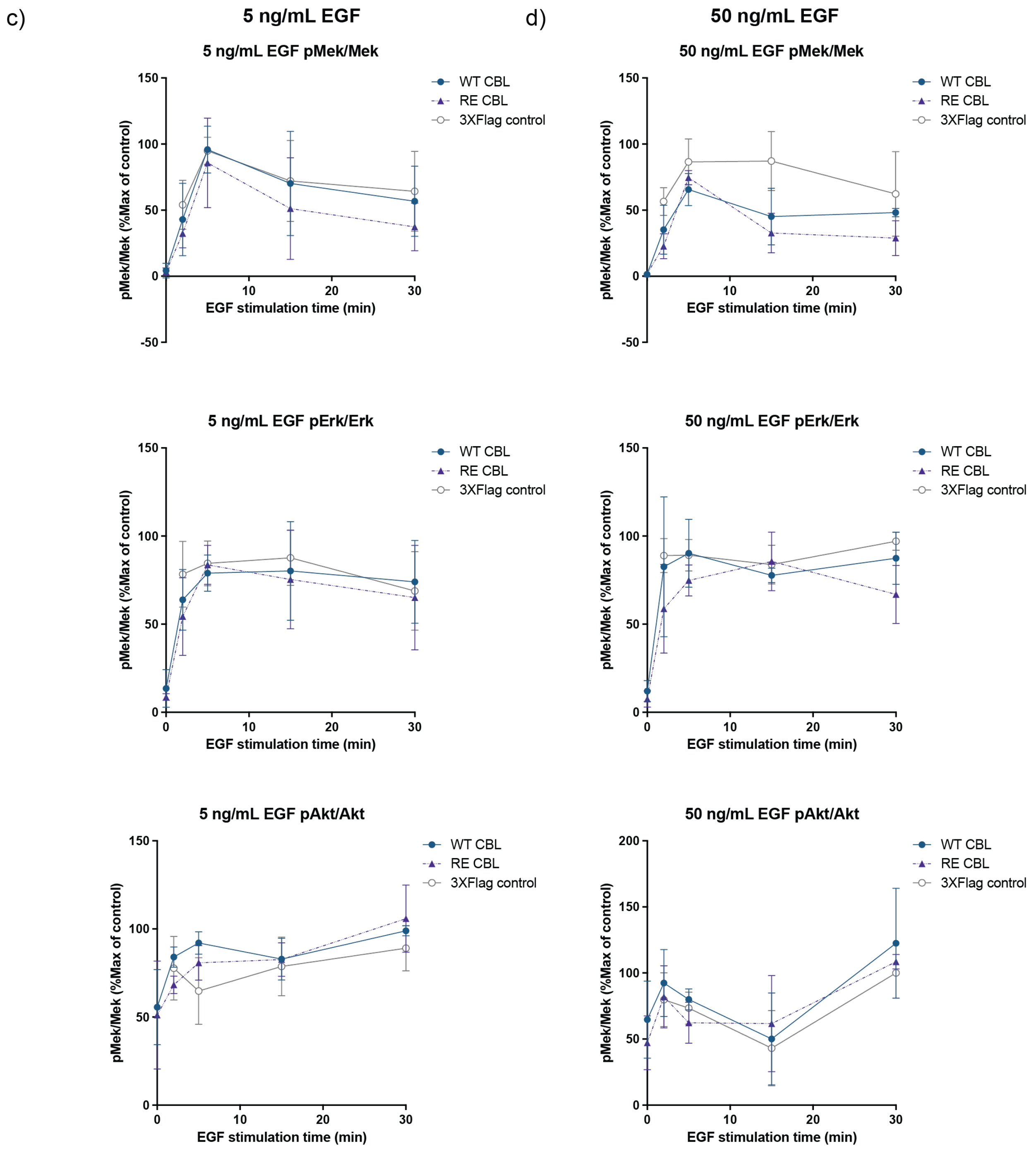


**Supplementary Figure 5: Assessing signalling downstream of EGFR in the presence of RE CBL under high and low EGF concentrations**

HeLa Flp-In T-REx cells expressing C terminally 3XFLAG-tagged wildtype (WT) or RE CBL or FLAG control were seeded and induced for 8 hours in full serum and then changed to serum starved conditions in DMEM with 1 µg/mL tetracycline for an addition 16 hours. Cells were then all treated with **a)** 5 ng/mL or **b)** 50 ng/mL EGF at 37°C over a time course. Cells were washed with PBS and lysed. Quantified lysates were prepared to run out samples of equal protein level across conditions for immunoblotting for EGFR, CBL (FLAG) and downstream signalling proteins including: active MEK1/2 (Phospho-MEK1/2 (Ser217/221); pMEK – from **Figure 5c**), active Erk1/2 (pErk1/2 Thr202, Tyr204; pErk) and active Akt (Phospho-Akt (Ser473); pAkt). Blots were subsequently stripped and re-probed for total MEK, Erk and Akt respectively. Figures shown are representative of 4 biological replicates for **a)** 5 ng/mL EGF treatment and 3 biological replicates for **b)** 50 ng/mL EGF treatment. Phosphorylated and total protein levels were quantified for MEK, Erk and Akt. Phosphorylated proteins were normalized to the total protein levels and overall conditions were normalized to the maximum signal in the control over time course for each experiment. In **c)** and **d)**, the plots for quantification of the pMEK1/2, pErk and pAkt signal are shown for **c)** 5 ng/mL EGF blots and **d)** 50 ng/mL EGF blots.

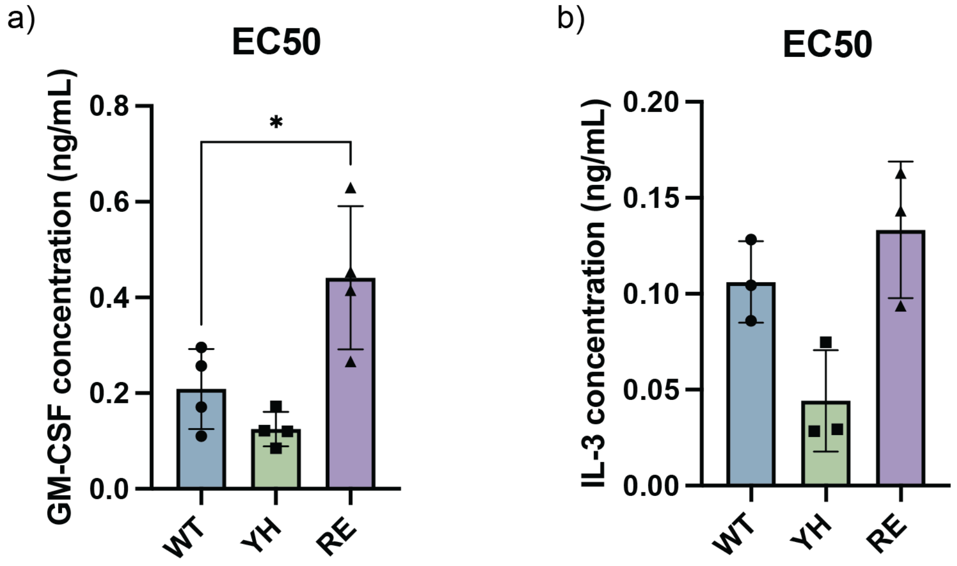


**Supplementary 6: 32D cell cytokine growth response EC50 values**

Plots of corresponding EC50 values for data in **Figure 6a** and **6b** for cells grown in **a)** GM-CSF or **b)** IL-3. EC50 values calculated for independent biological replicates and plotted with the points representing individual experiments and mean EC50 values represented by the bar and error bars showing the standard deviation. EC50 values were compared with by one-way ANOVA with Dunnett correction to compare WT CBL expressing cells and each other line (* indicates $p < 0.05$).

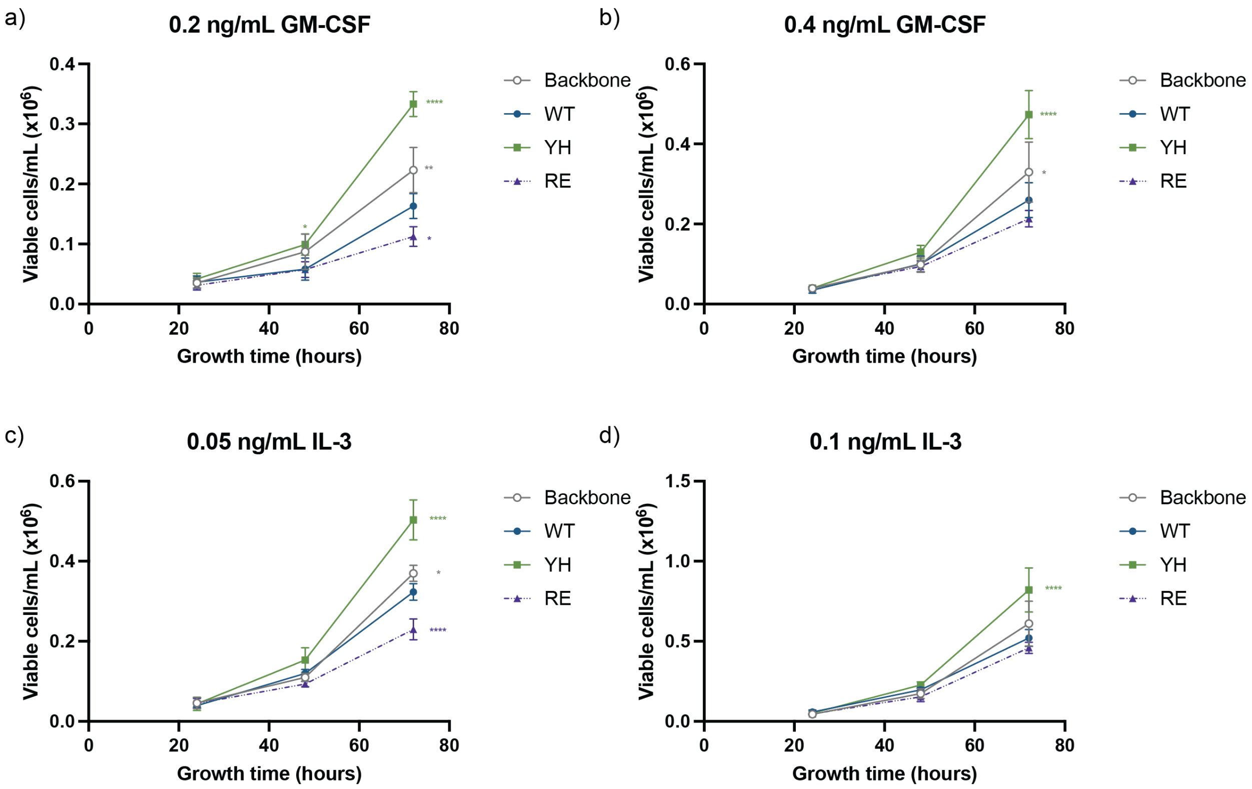


**Supplementary 7: 32D cell counts over time**

Cell counts of cultures grown for the immunoblot experiments in **Figure 6** (GM-CSF) assessed from 500 μL samples taken from 20 mL cultures every 24 hours from the initial set up. Results with cells grown with **a)** 0.2 ng/mL GM-CSF, **b)** 0.4 ng/mL GM-CSF, **c)** 0.05 ng/mL IL-3 and **d)** 0.1 ng/mL IL-3. The growth curves show the average of triplicate experiments at each time for each cell line with significance assessed by two-way ANOVA with Dunnett correction, and significance values shown relative to cells expressing WT CBL and the other mutants or backbone (* $p<0.05$, ** $p<0.01$, *** $p<0.001$, **** $p<0.0001$).

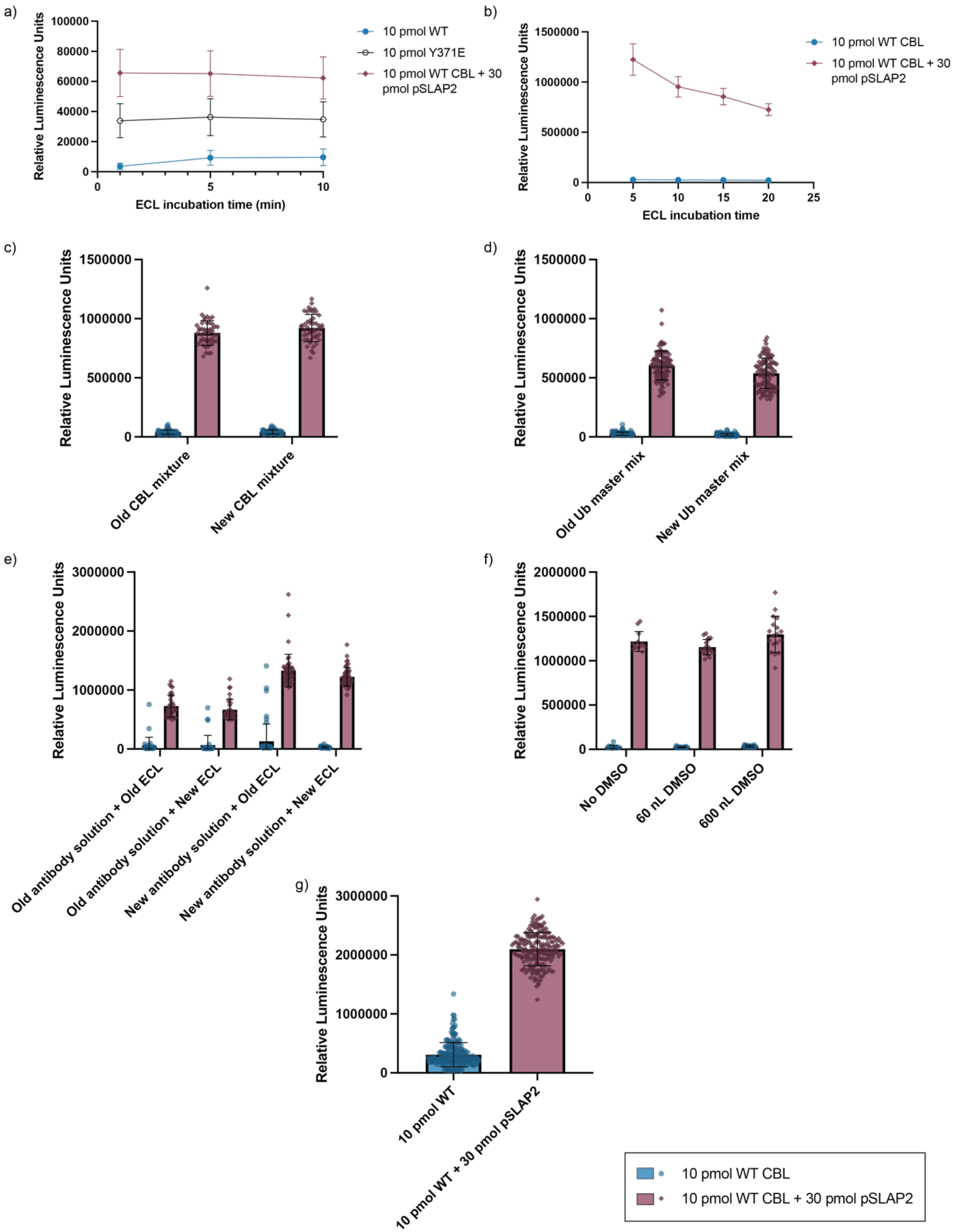


**Supplementary 8: Small Molecule Screen Optimization**

To determine the ideal timing between the addition of the enhanced chemiluminescent (ECL) reagent and the luminescence read of the plate on the Synergy Neo plate reader, a plate was setup with alternating duplicate columns of WT CBL, Y371E CBL and WT CBL with pSLAP2. Half of the plate was read in a manner where the ECL was added by the Synergy Neo plate reader well by well, and the signal was acquired immediately after each addition (inject/read). Data not shown due to low signal. **a)** The other half of the plate had the ECL added by the Multidrop Combi and then read right after the ECL was incubated, and the plate was read 1, 5 and 10 min after the ECL addition. Outlier wells due to technical issues in the assay set up were excluded in the plot showing signal over time. The point represents the mean signal from each condition at the indicated times, with error bars representing the standard deviation. WT data is shown in blue, signal with Y371E in black and signal with WT CBL with pSLAP2 in red. The addition of ECL by the Multidrop Combi was the selected parameter for the assay. Throughout further optimization, the 5 minute incubation time was reconfirmed in optimization as for example in **b)** where reagent stability was also assessed and signals for "new reagents" only are shown. **c-e)** To ensure consistency in signals throughout screening day, the stability of the positive (CBL+pSLAP2) and negative (CBL alone) controls were assessed with "new" reagent mixtures made just before addition to the assay plate, or "old" mixtures, that were left to sit on ice three to three and a half hours before the addition to the assay plate. All data are grouped by solution condition and the values for CBL (blue bar and circular points for individual wells) shown and then CBL + pSLAP2 beside (red bar and diamond shaped points for individual wells). For all graphs, bars represent the mean of the wells for each condition, and the error bars show the standard deviation. Individual graphs show assessment of: **c)** the stability of the CBL and CBL+pSLAP2 mixtures, **d)** the stability of the ubiquitination master mix, **e)** the antibody solution and enhanced chemiluminescent substrate (ECL) were tested together. **f)** To determine the effect of the compound solvent, DMSO, on the reactions, the controls were run with the DMSO that would be. The graph shows the comparison of the CBL and CBL+pSLAP2 signals with no DMSO, 60 nL DMSO or 600 nL DMSO (library concentration). **g)** Shows the results from final control plate before the screen was run. For each graph, bars alternate with the CBL signal and CBL+pSLAP2 signals of the relative luminescence detected by the plate reader. The legend for all graphs in this figure is shown on the bottom right.

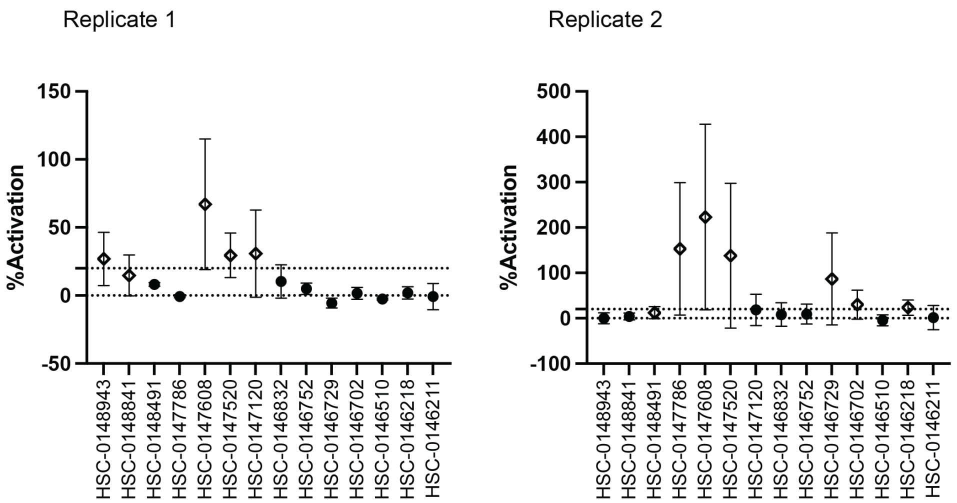


**Supplementary Figure 9: Screen secondary validation**

Two hit validation plate experiments were run with hit compounds assessed in triplicate wells of an individual plate that also included CBL and CBL with pSLAP2 (both with DMSO) as the negative and positive controls respectively. The results from both replicate experiments are plotted with the mean %activation of the three technical wells for each compound shown with the standard deviation indicated. The individual compounds are shown on the x-axis. Compounds that had at least 2 of 3 wells with a 20% activation were considered confirmed hits and are shown as open points. Compounds were considered confirmed hits if they met these criteria in either experiment.

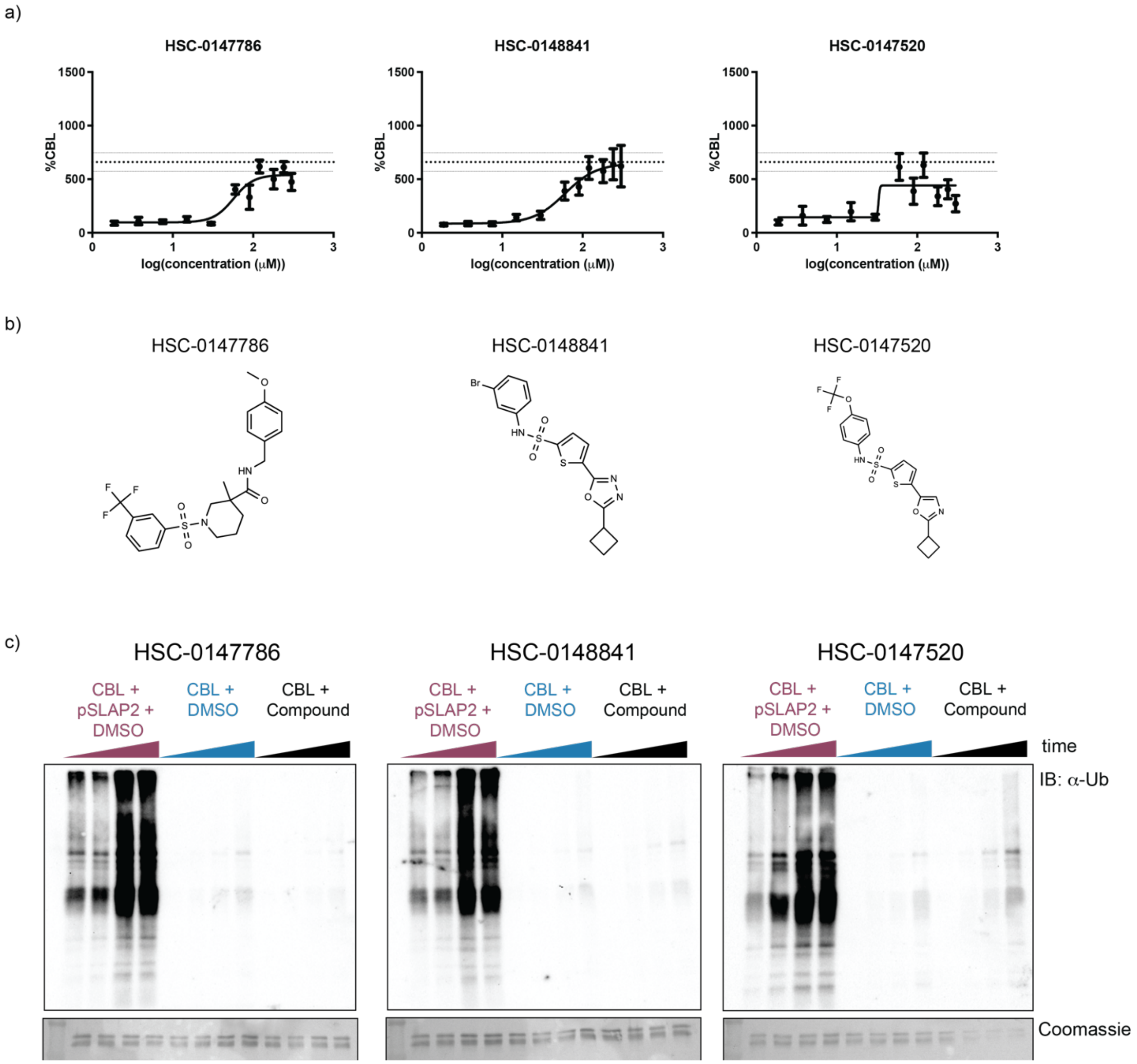


**Supplementary Figure 10: Peptidomimetic hits related to HSC-0147608**
**a)** Dose response curves for additional Peptidomimetic hit compounds displaying some degree of dose dependency. This data encompasses three independent experiments, where each concentration had duplicate wells per plate. Each plate also included wells of the negative control, CBL alone, and the positive control, CBL + pSLAP2. The log of the concentrations are plotted on the x-axis. To more accurately combine the three replicates, the relative luminescent values were normalized to the average CBL signal for the respective plates and then multiplied by 100 giving the %CBL signal. The mean of 6 values was plotted with the standard deviations against the log of the concentrations on the y-axis. The horizontal lines indicate the average %CBL for the positive control, CBL + pSLAP2, with the upper and lower lines showing the boundaries of the standard deviation. **b)** Compound structures for corresponding compounds in **a)** and **c)**. **c)** *In vitro* ubiquitination reactions were run over a time course with 1200 pmol (48 μM) of compounds with HSC-0147786, HSC-0148841 and HSC-0147520. Reactions master mixes were setup and then separated into individual tubes with 25 μL reactions. Reactions were stopped by the addition of sample buffer and immediately boiled after 30, 60, 90 or 120 minutes

of incubation. The amount of ubiquitination was visualized by immunoblot with probing for ubiquitin. Each blot included samples of positive control of CBL with pSLAP2 (and DMSO), CBL and no activating component with DMSO and CBL with the corresponding compounds of the labels shown above the blots. Timing of the reactions are from low (30 min) to high (120 min) for each condition as indicated. Lower images show the Coomassie stained membranes with CBL bands visible (SLAP2 was run off the gels, bands show CBL protein).

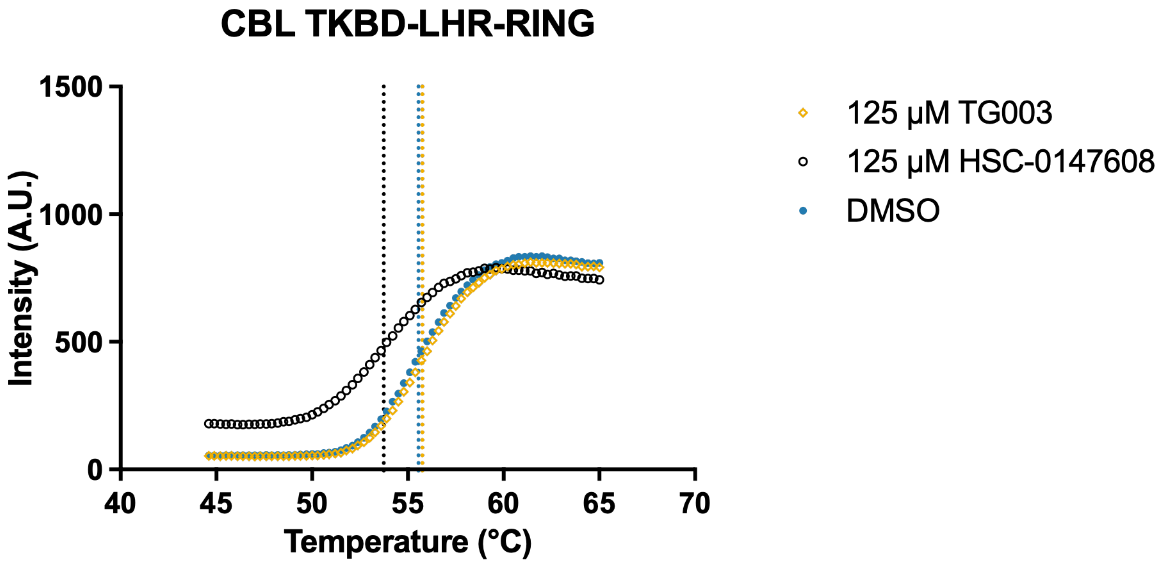


**Supplementary Figure 11: CBL Thermodenaturation is not impacted by an unrelated small molecule**

Thermodenaturation of $CBL_{2-436}$ over a temperature gradient in the presence of DMSO, 125 µM HSC-0147608 or 125 µM TG003 (CLK1 inhibitor). Vertical lines indicate the temperature of aggregation for 125 µM (53.74°C) HSC-0147608, DMSO (5%; 55.55°C), 125 µM TG003 (55.75°C).

a)

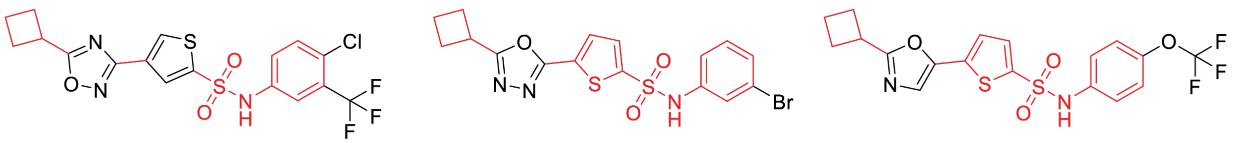


HSC-0147608 HSC-0148841 HSC-0147520

b)

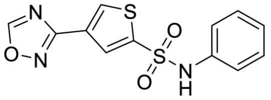


4 compounds (0.13% of library)

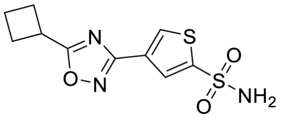


9 compounds (0.3% of library)

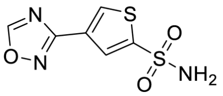


16 compounds (0.53% of library)

**Supplementary Figure 12: Structure comparison of Peptidomimetic hit compounds**
**a)** Comparison of the three related hit compounds. Red highlighted regions indicate conserved structures. **b)** Frequency of HSC-0147608 substructures in the 3000 compound library.

**Supplementary Table 1: Z' Statistics for Small Molecule Screen**

| Plate | Z′-factor (controls) | Positive control mean (n = number of wells) (RLU) | Negative control mean (n = number of wells) (RLU) |
|---|---|---|---|
| **Primary Screen** | | | |
| PM-384-1 | 0.36 | 2.28E+06 ± 410E+03 (n = 16) | 128E+03 ± 46.2E+03 (n = 32) |
| PM-384-2 | 0.29 | 1.30E+06 ± 255E+03 (n = 32) | 57.6E+03 ± 40.5E+03 (n = 32) |
| PM-384-3 | 0.31 | 2.57E+06 ± 390E+03 (n = 32) | 336E+03 ± 123E+03 (n = 32) |
| PM-384-4 | 0.39 | 2.91E+06 ± 281E+03 (n = 32) | 488E+03± 214E+03 (n = 32) |
| PM-384-5 | 0.37 | 2.61E+06 ± 235E+03 (n = 32) | 525E+03± 201E+03 (n = 32) |
| PM-384-6 | 0.46 | 2.90E+06 ± 296E+03 (n = 32) | 465E+03 ± 141E+03 (n = 32) |
| PM-384-7 | 0.42 | 2.89E+06 ± 275E+03 (n = 32) | 564E+03 ± 179E+03 (n = 32) |
| PM-384-8 | 0.31 | 2.69E+06 ± 289E+03 (n = 32) | 558E+03± 203E+03 (n = 32) |
| PM-384-9 | 0.31 | 2.57E+06 ± 375E+03 (n = 32) | 328E+03 ± 143E+03 (n = 32) |
| PM-384-10 | 0.22 | 2.41E+06 ± 294E+03 (n = 32) | 514E+03± 199E+03 (n = 31) |
| **Hit Validation** | | | |
| Hit confirmation #1 | 0.3 | 1.76E+06 ± 240E+03 (n = 16) | 229E+03 ± 117E+03 (n = 112) |
| Hit confirmation #2 | -0.08 | 870E+03 ± 188E+03 (n = 53) | 199E+03 ± 54.3E+03 (n = 42) |
| **Dose Response** | | | |
| Replicate #1 | 0.17 | 2.69E+06 ± 401E+03 (n = 48) | 426E+03 ± 228E+03 (n = 140) |
| Replicate #2 | 0.38 | 3.14E+06 ± 370E+03 (n = 48) | 460E+03 ± 184E+03 (n = 140) |
| Replicate #3 | 0.41 | 2.95E+06 ± 332E+03 (n = 48) | 42E+03 ± 228E+03 (n = 140) |

### Supplementary Methods

#### Co-immunoprecipitation

HeLa Flp-In T-REx cells stably transfected to express carboxy-terminal 3XFLAG tagged CBL (wildtype or RE) were seeded on 10 cm dishes. Cells were induced the next day in DMEM with 1% penicillin/streptomycin and 10% tetracycline-free FBS with 1 μg/mL tetracycline for 32 hours. Media was subsequently changed to DMEM alone with 1 μg/mL tetracycline for an additional 16 hours. Induction media was aspirated, and cells were treated with EGF (1 ng/mL, 2.5 ng/mL, 5 ng/mL, 10 ng/mL, 50 ng/mL or 100 ng/mL diluted in DMEM) in 3 mL of media per plate. Cells were incubated at 37°C for 2 minutes. Treatments were conducted in groups by EGF concentration, with all cell lines treated concurrently. After the incubation, cells were immediately transferred to ice and washed with cold PBS. Cells were lysed in 800 μL of PLC lysis buffer (50 mM HEPES pH 7.5, 150 mM NaCl, 1.5 mM $MgCl_2$, 1 mM EDTA (pH 8), 10% glycerol, 1% Triton X-100, supplemented with cOmplete protease inhibitor table (Roche #05056489001), $Na_3VO_4$, and 10 mM NaF). Cell lysates were incubated on ice for at least 30 minutes and then sonicated for 5 seconds. Lysates were cleared by centrifugation at 20817 rcf for 15 minutes at 4°C. Cleared lysates were transferred to new tubes and lysates were quantified using the Pierce BCA Protein Assay Kit. Lysates were used concurrently in FLAG (CBL) and EGFR immunoprecipitation experiments. For the FLAG pull down, 20 μL of packed resin (ANTI-FLAG® M2 Affinity Gel, Millipore #A2220) per sample was washed three times in lysis buffer with spins at 5000 rcf for 30 seconds between washes. 500 μg of lysates for each sample was added to the beads, with extra lysis buffer added to equalize the volumes. Samples were incubated for 4 hours at 4°C with gentle rocking. EGFR immunoprecipitation samples were prepared with 500 μg of lysates incubated with 2 μg of anti-EGFR antibody (EGFR Polyclonal

Antibody, ThermoFisher Scientific #PA1-1110) for 2 hours at 4°C with gentle rocking. 20 µL of protein A Sepharose (ThermoFisher Scientific #101041) per sample was washed three times as outlined above. The Sepharose was then resuspended to add 100 µL to each EGFR immunoprecipitation after the initial 2 hour incubation, and the samples were incubated for an additional 2 hours at 4°C with gentle rocking. After the incubation, all immunoprecipitation samples were washed three times with 500 µL of lysis buffer with centrifugation at 5000 rcf for 30 seconds at 4°C between each wash. After the final wash, samples were spun an additional time and resin was dried through aspiration of any residual buffer. Resin was resuspended in 70 µL of 1X sample buffer (62.5 mM Tris pH 6.8, 2% SDS, 10% glycerol, 0.3575 M β-mercaptoethanol, bromophenol blue), mixed, boiled for 10 minutes and then spun down for 1 minute at 20800 rcf. Lysate samples of 30-40 µg were also prepared for immunoblotting with 6X sample buffer followed by boiling for 10 minutes and centrifugation. Samples were stored at -20°C until immunoblot. For the immunoprecipitation samples, 32 µL of the samples were resolved on per gel. For quantification, background adjusted volume measurements, were obtained for EGFR and FLAG for each experiment. For the FLAG immunoprecipitation, EGFR intensity values were normalized to FLAG, and for the EGFR immunoprecipitation, FLAG intensity values were normalized to EGFR for each condition. The normalized signals were computed as a percent maximum signal ratio in each experiment and the normalized values across three independent biological replicates were plotted. Significance values are shown only between WT and RE not within the same cell line across time on the graph.

**CBL level comparison with MG132 treatment**

HeLa Flp-In T-REx cells expressing carboxy-terminally tagged FLAG-MiniTurboID biotin ligase with wildtype or RE CBL were induced with tetracycline in full media for 32 hours (or left uninduced) and changed to starvation conditions with tetracycline and indicated concentrations of MG132 (Millipore Sigma #474790; prepared in DMSO and diluted in media, 0 μM condition had DMSO) 16 hours before lysis. HeLa Flp-In T-REx cells expressing carboxy terminally tagged 3XFLAG tagged wildtype or RE CBL clonal lines that were induced in full serum media for 8 hours with or without tetracycline and then switched to DMEM only starved conditions with increasing concentrations of MG132. Cells were lysed after 16 hours. Following the incubation with MG132, cells were cooled on ice and washed twice with cold PBS. Cells were lysed in 100 μL of modified RIPA (50 mM Tris pH 7.5, 150 mM NaCl, 0.4% SDS, 1% NP-40, 1.5 mM MgCl2, 1 mM EGTA, supplemented with 10 μL of benzonase in 10 mL, mini cOmplete protease inhibitor tablet, and Na3VO4). Cell lysates were incubated on ice for at least 30 minutes and then cleared by centrifugation at 20817 rcf for 15 minutes at 4°C. Cleared lysates were transferred to new tubes and lysates were quantified using the Pierce BCA Protein Assay Kit. 30 μg lysate samples were prepared to run on immunoblots. FLAG (CBL) and tubulin levels were quantified. FLAG signals were normalized to tubulin levels and the normalized signals were computed as a percent of the wildtype signal with tetracycline only in each experiment and the normalized values across three independent biological replicates were plotted. Significance values are shown only between wildtype and RE CBL expressing cells, not within the same cell line on the graph.

**Surface level protein capture biotinylation assay**

HeLa Flp-In T-REx cells stably transfected to express carboxy terminal 3XFLAG tagged CBL (wildtype or RE) or the FLAG control were seeded on 10 cm dishes with one plate seeded per time point per cell line. Cells were induced the next day in DMEM with 1% penicillin/streptomycin and 10% tetracycline-free FBS with 1 µg/mL tetracycline for 8 hours and then media was changed to DMEM with 1 µg/mL tetracycline for 16 hour starvation. Cells were treated with 5 or 50 ng/mL EGF in 5 mL of DMEM and incubated over a time course at 37°C. Following the time point, cells were cooled on ice and washed twice with cold PBS. Cells were then incubated with 0.2 mg/mL EZ-Link™ Sulfo-NHS-SS-Biotin (Thermo Fisher Scientific #21331) made up fresh in biotinylation buffer (154 mM NaCl, 10 mM HEPES, 3 mM KCl, 1 mM $MgCl_2$, 0.1 mM $CaCl_2$, 10 mM glucose, pH 7.6) for 1 hour at 4°C to label surface level proteins. After labeling, cells were washed twice with cold PBS and blocked for 5 minutes in DMEM with 10% tetracycline-free FBS, 1% penicillin/streptomycin, 100 mM glycine at 4°C. Cells were then washed again two times with cold PBS and lysed in 500 µL of PLC lysis buffer (50 mM HEPES pH 7.5, 150 mM NaCl, 1.5 mM $MgCl_2$, 1 mM EDTA (pH 8), 10% glycerol, 1% Triton X-100, supplemented with cOmplete protease inhibitor table, $Na_3VO_4$, and 10 mM NaF). Cell lysates were incubated on ice for at least 30 minutes and then cleared by centrifugation at 20817 rcf for 15 minutes at 4°C. Cleared lysates were transferred to new tubes and lysates were quantified using the Pierce BCA Protein Assay Kit. 30 µL of streptavidin agarose (Thermo Fisher Scientific #20353) per sample was washed as a bulk solution for the total number of samples. The agarose resin was washed three times with the lysis buffer with 2 minute spins at 2000 rcf at 4°C between each wash. Resin was resuspended in lysis buffer to aliquot 100 µL of washed resin to new tubes and 350 µg of lysate was added. Lysis buffer was then added to equalize the volumes. Samples were incubated overnight at 4°C with gentle rocking. 20 μg of lysate was also prepared for immunoblot in sample buffer, with boiling for 10 min and spin down at 208700 rcf for 1 minute before storing. The next day, pull down samples were washed 4 times with lysis buffer with 2 minute spins at 2000 rcf at 4°C between each wash. After the final wash, samples were spun an additional time and resin was dried through aspiration of any residual buffer. Resin was resuspended in 70 μL of 1X sample buffer (62.5 mM Tris pH 6.8, 2% SDS, 10% glycerol, 0.3575 M β-mercaptoethanol, + bromophenol blue), mixed, boiled for 10 minutes and then spun down for 1 minute at 20800 rcf. Samples were assessed by immunoblot as described for the immunoprecipitation experiment, with 32 μL of the pull down run on one gel. Membranes were cut and probed as indicated. For the surface level protein pull down blots, EGFR and transferrin levels were quantified. EGFR signals were normalized to transferrin levels, and the normalized signals were computed as a percent of the wildtype signal at 0 minutes of EGF in each experiment. Lysate blots were quantified for EGFR and tubulin. EGFR levels were normalized to tubulin level and then to the signal for wildtype CBL at 0 ng/mL EGF in each experiment. In both cases, the normalized values across three independent biological replicates were plotted. Significance values are shown only between WT and RE CBL expressing cells across time on the graph.

**Time course of EGFR signalling activation**

HeLa Flp-In T-REx cells stably transfected to express carboxy terminal 3XFLAG tagged CBL (wildtype or RE) or the FLAG control were seeded on 6 well plates with one well seeded per time point per cell line (30, 15, 5, 2 minutes and no EGF). Cells were induced the next day in DMEM with 1% penicillin/streptomycin and 10% tetracycline-free FBS with 1 μg/mL tetracycline for 8 hours, and then media was changed to DMEM with 1 μg/mL tetracycline for 16 hour starvation. Cells were treated with 5 or 50 ng/mL EGF in 5 mL of DMEM and incubated over a time course at 37°C. Following the time point, cells were cooled on ice and washed twice with cold PBS. Cells were lysed in 80 μL of PLC lysis buffer (50 mM HEPES pH 7.5, 150 mM NaCl, 1.5 mM $MgCl_2$, 1 mM EDTA (pH 8), 10% glycerol, 1% Triton X-100, supplemented with cOmplete protease inhibitor tablet (Roche), $Na_3VO_4$, and 10 mM NaF). Cell lysates were incubated on ice for at least 30 minutes and then cleared by centrifugation at 20817 rcf for 15 minutes at 4°C. Cleared lysates were transferred to new tubes and lysates were quantified using the Pierce BCA Protein Assay Kit (Thermo). 30 μg lysate samples were prepared to run on multiple immunoblots. Phosphorylated proteins (Akt, Erk1/2 and Mek1/2) were assessed and then membranes were stripped in harsh stripping buffer (2% SDS, 62.5 mM Tris HCl pH 6.8, 114.4 mM β-mercaptoethanol) warmed to 50°C with agitation for 45 minutes. Stripped membranes were then washed in water and then TBST before being blocked for 1 hour in 1% fish gel in TBST and re-probed for the corresponding total protein. Phosphorylated and total levels of Akt, Mek1/2 and Erk1/2 were quantified. Signals were normalized with phosphorylated protein levels relative to total and the normalized signals were computed as a percent of the maximum signal observed for the control cell line in each experiment. The normalized values across three independent biological replicates were plotted.

**CBL modification time course immunoprecipitation**

HeLa Flp-In T-REx cells stably transfected to express carboxy terminal 3XFLAG tagged CBL (wildtype or RE) were seeded on 10 cm plates with one plate seeded per time point per cell line (30, 15, 5, 2 minutes and no EGF). Cells were induced the next day in DMEM with 1%

penicillin/streptomycin and 10% tetracycline-free FBS with 1 µg/mL tetracycline for 8 hours and then media was changed to DMEM with 1 µg/mL tetracycline for 16 hour starvation. Cells were treated with 5 ng/mL EGF in 5-6 mL of DMEM and incubated over a time course at 37°C (from longer to shorter time points). Following each time point, plates were cooled on ice and washed twice with cold PBS. Cells were lysed in 700 µL of RIPA lysis buffer (1% Triton X-100, 50 mM Tris pH 7.5, 150 mM NaCl, 1 mM EDTA, 0.1% SDS, 1.5 mM $MgCl_2$, 1% sodium deoxycholate with cOmplete protease inhibitor tablet (Roche) and $Na_3VO_4$, as well as 1µL/mL benzonase nuclease in lysis). Following all time points, lysates were incubated in 1.5 mL tubes rotating at 4°C and then sonicated for 10s with 40% amplitude. Samples were then spun for 15 min at 21300 rcf at 4°C and cleared lysate transferred to fresh tubes. Samples were quantified using the Pierce BCA Protein Assay Kit. For the FLAG pull down, 20 µL of packed resin (ANTI-FLAG® M2 Affinity Gel A2220, Millipore) per sample was washed three times in lysis buffer with spins at 5000 rcf for 30 seconds between washes. 400 µg of lysates for each sample was added to the beads, with extra lysis buffer added to equalize the volumes. Samples were incubated for 4 hours at 4°C with gentle rocking. Immunoprecipitation samples were washed and prepared as per the co-immunoprecipitation with RIPA buffer, and 20 µg lysate samples were prepared. Immunoprecipitation samples were equally split into 3 gels for immunoblot. For immunoprecipitation blots, pY371 or ubiquitin signals were quantified and plotted as a percentage of the maximum signal with wildtype CBL in the experiment. For lysate samples, the levels of the upper bands of EPS15 were quantified relative to the total EPS15 levels expressed as %Modified EPS15. Significance values are shown only between WT and RE not within the same cell line across time on the graph.

**32D cell proliferation and immunoblots**

32D cells were washed twice with PBS with spins at 0.1 rcf for 7 min and resuspended in induction media (RPMI with 10% tetracycline-free FBS with 500 ng/mL doxycycline with indicated concentrations of IL-3 or GM-CSF). 500 µL of cells were used to count using a ViCell XR cell counter (Beckman-Coulter). $0.3 \times 10^6$ cells for each 32D cell line were seeded to have 20 mL of induction media total. 24, 48 and 72 hours after seeding, cells were resuspended and 500 µL was removed to count. At 72 hours, after the sample was removed to count, cells were spun at 1000 rpm for 5 minutes at 4°C and washed with 10 mL of cold PBS. Cells were spun again and then resuspended in 1 mL of cold PBS and transferred to a 1.5 mL tube. Cells were spun again and PBS aspirated. Cell pellets were resuspended in boiling Laemmli buffer (0.0625 M Tris pH 6.8, 2% SDS and 10% glycerol). Lysates were sonicated for 5-10s and then boiled for 5 minutes. Lysates were cleared by centrifugation at 208000 rcf for 15-20 minutes before transferring to fresh tubes. Lysates were quantified using the Pierce BCA Protein Assay Kit. 40µg lysate samples were prepared with 6X SDS sample buffer to run on multiple immunoblots (phospho-Lyn and total Lyn assessed on independent blots from the same lysates). Lyn or phospho-Lyn levels were and normalized to GAPDH levels. The normalized signals were computed as a percent of the wildtype signal in each experiment and the normalized values across three independent biological replicates were plotted. Cell counts over time were analyzed by two-way ANOVA with Dunnett correction to compare mutants and backbone to wildtype in GraphPad Prism 10.

**Immunofluorescence**

HeLa Flp-In T-REx cells stably transfected to express the C terminal FLAG-MiniTurbo fusion with WT or RE CBL, or the MiniTurbo control line were seeded on glass coverslips in 24 well plates. The next day, cells were induced with 1 μg/mL tetracycline (in the biotin labeling experiment, MiniTurbo control cells were induced with 5 ng/mL tetracycline, while in the EGF treatment experiment, the control cells were also induced with 1 μg/mL tetracycline) in DMEM with 1% penicillin/streptomycin, with 10% FBS (biotin depleted, tetracycline-free). Cells were induced for 32 hours before media was changed to DMEM alone with fresh tetracycline. Cells were treated as described in the figure captions with media changes to introduce conditions with either 50 μM biotin or 100 ng/mL EGF as outlined. Time course incubations were conducted with cells incubated at 37°C. Following cell treatment, media was aspirated, and coverslips were washed 2-3 times with PBS (with $Ca^{2+}/Mg^{2+}$). Cells were fixed in 2% paraformaldehyde for 10 minutes and washed 2-3 times with PBS (with $Ca^{2+}/Mg^{2+}$). Washes were two times for the EGF treatment and three times for the biotin treatment. Paraformaldehyde fixation was quenched with 100 mM glycine in PBS incubated for 10 min. Coverslips were washed 2 times with PBS (with $Ca^{2+}/Mg^{2+}$). Cells were permeabilized with 0.1% Triton X-100 in PBS (with $Ca^{2+}/Mg^{2+}$) for 10 minutes followed by 3 washes with PBS (with $Ca^{2+}/Mg^{2+}$). Cells were blocked in 5% donkey serum in PBS-T (0.1% tween) for 30 minutes at room temperature. Anti-FLAG M2 (Sigma # F3165) antibody was diluted 1 in 2000 with 1:50 anti-EGFR antibody (Cell Signaling Technology # D38B1) in blocking buffer for the EGF treated cells. Streptavidin labeling was done with the secondary and no EGFR costain. Primary antibody was applied overnight with coverslips incubated at 4°C in a humidified box. The next day, coverslips were washed 3 times with PBS-T (0.05% Triton X-100) for 10 minutes per wash with agitation. Secondary antibody

(1:500 Alexa488 donkey anti-mouse IgG (Invitrogen #A21202), 1:500 Cy3 donkey anti-rabbit IgG (Jackson ImmunoResearch #711-165-152) or 1:2500 streptavidin-Alexa647 (Invitrogen #S32357)) was diluted in blocking buffer and incubated on coverslips in a humidified box at 37°C for 30 minutes. Cover slips were washed 3 times with PBS-T (0.05% Triton X-100) for 10 minutes per wash with agitation. Coverslips were rinsed in MilliQ water and then mounted on slides using Dako Fluorescent mounting media (Agilent #S3023) and allowed to set at least overnight. Images were acquired with 0.3 µm Z stack using a spinning disk confocal (Olympus IX81) on 60X oil objective with a Hamamatsu C9100-13 EM-CCD camera. Images were processed with iterative deconvolution on Volocity software using 100% confidence and 30 (streptavidin images) or 20 (EGFR co-stain) iterations. Snapshots of individual planes at similar regions of the cell are shown. Experiments were conducted independently at least twice.

**Protein purification**

$CBL_{2\text{-}436}$ was overexpressed in *Escherichia coli* BL21 (1 L Luria Bertani (LB) media supplemented with 100 µg/ml ampicillin per culture, grown to OD600 of 0.5-0.8, incubated overnight at 18°C with 0.5 mM IPTG induction; for the primary screen, two independent protein preparations of 6 L and 8 L of culture were combined). Cultures were pelleted and stored at -80°C. Bacterial cell pellets were lysed in 50 mM HEPES (pH 7.5), 150 mM NaCl, 1.5 mM $MgCl_2$, 1 mM EDTA pH 8.0, 10% glycerol, 1% NP-40 supplemented with benzonase nuclease (Millipore #70746-3) and cOmplete protease inhibitor tablet (Roche #05056489001). Lysates were subsequently supplemented with 0.5 mM PMSF following sonification. Lysate was cleared by centrifugation and supernatant was mixed with glutathione Sepharose 4B (GE Healthcare #17-0756) for 3 hours by nutating at 4°C. Resin was washed 3x25mL with NP-40 wash (20 mM

HEPES, 150 mM NaCl, 5% glycerol, 0.1% NP-40, with 0.5 mM PMSF, 5 mM beta-mercaptoethanol (βME)) and 3x25 mL washed with PBS wash (PBS with 5 mM βME) on a gravity column. Resin was incubated with thrombin protease (Sigma #T4648) in PBS (5 mM βME) and CBL protein eluted with PBS (5 mM βME). Following combination of the two purified protein purifications, the concentration was determined by Bradford assay (BioRad #5000006). Protein was aliquoted and then flash frozen in liquid nitrogen and stored at -80°C. The same protein preparations were used for the primary screen and secondary validation experiments. New preparations were completed for the dose response and subsequent validation experiments where several smaller scale purifications were conducted. Phosphorylated CBL protein was produced in the same manner except expressed in *Escherichia coli* TKB1 bacteria as described below for SLAP2, followed by purification in CBL buffers. For StarGazer-2, additional purification of the CBL protein was conducted by running on a Hi Load Superdex 16/600 200 pg column (Cytiva #28-98-93-35) using an AKTA pure FPLC equilibrated with PBS (5 mM βME) and isolating fractions containing the expected band size of approximately 49 kDa through assessment of collected fractions by SDS-PAGE.

Trx-His(6)-hSLAP2 WT was expressed in *Escherichia coli* TKB1 bacteria (12x100 mL cultures grown in LB supplemented with 50 μg/ml ampicillin and 12.5 μg/ml tetracycline per culture). Cultures were grown to an OD600 of 0.4-1 and induced with 0.5 mM IPTG overnight at 18°C. Cultures were combined, spun down and incubated in TK induction media (per manufacturer's instructions – Aligent #200134) for 3 hours at 37°C followed by centrifugation. Pellets were stored at -80°C. The bacterial pellet was lysed with 50 mM HEPES pH 7.5, 0.5 M NaCl, 10% glycerol, 1% Nonidet-P40, 10 mM imidazole, 1.5 mM $MgCl_2$ with 5 mM β-mercaptoethanol, 10 mM NaF, 1 mM $Na_3VO_4$, cOmplete protease inhibitor tablet. The lysate was treated with benzonase nuclease and sonicated. Following sonication, lysate was spiked with 0.5 mM PMSF and 0.5 mM $Na_3VO_4$. Phospho-SLAP2 was isolated using Ni-NTA agarose resin (Qiagen) incubated at 4°C for 3 hours. Resin was washed on a gravity column 6x15 mL of wash buffer (50 mM HEPES pH7.5, 0.5 M NaCl, 2% glycerol, 20 mM imidazole with 5 mM β-mercaptoethanol, 0.5 mM $Na_3VO_4$). pSLAP2 was eluted in a series of washes with increasing imidazole concentrations (75 mM, 150 mM, 225 mM, and 2x 300 mM; made up in wash buffer). The Trx-His(6)-tag was cleaved with TEV with simultaneous dialysis against (25 mM HEPES pH 7.5, 0.4 M NaCl, 4% glycerol, 10 mM imidazole, 5 mM βME, 5 mM $MgCl_2$, 0.5 mM $Na_3VO_4$) overnight at 4°C using slide-A-lyzer dialysis cassette 10000 MWCO 3-12 mL (Thermo Fisher Scientific #66810). Protein was removed from the dialysis cassette, spun at 4000 rpm for 7 min to remove precipitate, and supernatant was passed slowly over the NiNTA resin to remove the cleaved tag. Protein concentration was determined by Bradford assay and final protein was stored at 4°C or flash frozen in liquid nitrogen for long term storage (pSLAP2 used in the screen was stored at 4°C).

Commercial E1 (LifeSensors #UB-0101-0050) was used in reactions for the primary and secondary screen. Follow up dose response assays were conducted using in lab purified Uba1. Uba1 was overexpressed in *Escherichia coli* BL21 (1 L Luria Bertani (LB) media supplemented with 50 µg/ml kanamycin per culture, grown to OD600 of 0.8-1, incubated overnight at 18°C with 0.5 mM IPTG induction. Cultures were pelleted and stored at -80°C. Bacterial cell pellets were lysed in 20 mM HEPES pH 7.5, 400 mM NaCl, 5 mM imidazole pH 7.5, 10% glycerol with 1 mM PMSF. Lysates were subsequently supplemented with 0.5 mM PMSF following sonification. Lysate was cleared by centrifugation and supernatant was mixed with Ni-NTA agarose resin (Qiagen) and incubated for 2-3 hours by nutating at 4°C. Resin was washed 6 times with 15mL wash buffer (20 mM HEPES pH 7.5, 1000 mM NaCl, 20 mM imidazole, 2% glycerol). Uba1 was eluted in a series of washes with increasing imidazole concentrations (75 mM, 150 mM, 225 mM, and 2x 300 mM, all made up in wash buffer). Protein solution was dialyzed overnight against 20 mM HEPES pH 7.5, 50 mM NaCl, 2% glycerol using a slide-A-lyzer dialysis cassette 10000 MWCO 3-12 mL (Thermo Fisher Scientific #66810). Protein quantification was assessed by Bradford assay.
